# SIMPLEs: a single-cell RNA sequencing imputation strategy preserving gene modules and cell clusters variation

**DOI:** 10.1101/2020.01.13.904649

**Authors:** Zhirui Hu, Songpeng Zu, Jun S. Liu

**Affiliations:** Harvard University, Department of Statistics, Cambridge, 02138, United States

## Abstract

A main challenge in analyzing single-cell RNA sequencing (scRNASeq) data is to reduce technical variations yet retain cell heterogeneity. Due to low mRNAs content per cell and molecule losses during the experiment (called “dropout”), the gene expression matrix has substantial zero read counts. Existing imputation methods either treat each cell or each gene identically and independently, which oversimplifies the gene correlation and cell type structure. We propose a statistical model-based approach, called SIMPLEs, which iteratively identifies correlated gene modules and cell clusters and imputes dropouts customized for individual gene module and cell type. Simultaneously, it quantifies the uncertainty of imputation and cell clustering. Optionally, SIMPLEs can integrate bulk RNASeq data for estimating dropout rates. In simulations, SIMPLEs performed significantly better than prevailing scRNASeq imputation methods by various metrics. By applying SIMPLEs to several real data sets, we discovered gene modules that can further classify subtypes of cells. Our imputations successfully recovered the expression trends of marker genes in stem cell differentiation and can discover putative pathways regulating biological processes.

## Introduction

Single cell RNA sequencing technology has been widely used in discovering subtypes of cells in the immune system^1-3^, the nervous system^4–6^, and different diseases^7^, etc., and identifying gene modules controlling various cellular processes, e.g. the developmental process^8,9^ or under different stimuli^10^. A typical single cell RNA sequencing data set has many zero entries, which can come from two sources: expression level below the measurement limit (“off” state) and technical “dropout“^11^. In order to impute the missing values caused by dropout, we need to distinguish technical zeros from the true biological “off” state. Previous methods usually pool information from similar cells to do imputation. For example, MAGIC defines a diffusion process on the affinity graph of cells for imputation^12^; for each of the highly probable dropout genes, scImpute^13^imputes the dropout values in one cell by learning from the same gene in other similar cells, in which the weights of other cells are determined by the genes not severely impacted by dropout. Similarly, VIPER uses sparse non-negative regression to progressively learn the local neighborhood cells and impute the gene expression based on these cells^14^. These methods often over-smooth the gene expression ignoring the cell-to-cell variations, despite that a main purpose of single cell experiments is to identify biological heterogeneity of cells. Moreover, based on different gene functional groups, distances between cells can be different. The aforementioned methods define the nearby cells averaging over all the genes without considering distinctions among genes.

Different from previous methods, we model the structure of gene correlations across similar cells and allow for different variability of the imputed values for each gene group. The aggregated effects across multiple correlated genes can distinguish dropouts from low expression even if the signal to noise ratio is low for each individual gene. This additional freedom of gene-group specific imputations preserves the stochasticity of gene expression observed in scRNASeq data. Our method, termed as **SI**ngle cell RNASeq i**MP**utation and cel**L**clust**E**ring) (SIMPLE), infers the probability of the dropout event for each zero entry, and imputes technical zeros while maintaining biological zeros at a low level. The imputation process depends on gene correlations within similar cell types, which is modeled by a few common gene modules, as well as the gene-specific dropout rate. Although the dropout rate can be estimated from the empirical distribution of gene expression in the scRNASeq, it can interfere with the estimation of the gene correlation structure, especially for lowly expressed genes. Bulk RNASeq data, which reveal the average gene expression across cells and provide an extra source of information on the dropout rate per gene, can also be incorporated into SIMPLE. We name such an extension for integrating bulk RNASeq data **SIMPLE-B** and refer to our toolbox including SIMPLE and SIMPLE-B as **SIMPLEs**. In addition to obtaining imputed expression matrix as previous methods usually do, SIMPLEs can output clusters of cells and gene modules that distinguish different subtypes of cells or groups of samples as a byproduct. Also, SIMPLEs can provide measures of uncertainty for imputed values, which can be incorporated into downstream analyses. For example, from multiple imputations, SIMPLEs can provide a consensus matrix of clustering membership indicating the uncertainty of the clustering results. A recent method SCRABBLE^15^ also uses the bulk RNASeq information, but only as a constraint to the mean gene expression of the scRNASeq data instead of a means of estimating dropout rates. Furthermore, it assumes similar gene expression in a few cell types and does not consider correlations among the genes. Another method SAVER^16^ takes advantage of gene-gene correlations for imputation, but does not model dropouts explicitly and treats each cell independently without pooling information from closely related cells. DEsingle^17^ can distinguish dropouts from biologically low expression using a zero-inflated model, but its goal is to detect differentially expressed genes.

By simulating the entire data set or adding more dropouts to a published scRNASeq data, we demonstrate the superior performances of SIMPLEs in gene expression imputation and cell clustering compared with prevailing methods for scRNASeq imputation. Then, we applied SIMPLEs to two real data sets: human embryonic stem cells differentiation and mouse preimplantation embryos. In both data sets, we discovered gene modules that can further classify subtypes of cells. Moreover, the imputed values of marker genes by SIMPLEs aligned well with the developmental stages of each cell, suggesting that SIMPLEs can be used to discover gene markers that regulate the developmental process. Finally, we manifest the scalability of SIMPLEs and its robustness on different parameters using a large scale data set of mouse immune cells from multiple tissues.

## Results

### Framework of SIMPLEs

Given the log-normalized data (e.g., logarithm of one plus RPKM, FPKM, or TPM), we model the gene expression within a cell type by a zero-inflated censored multivariate Gaussian distribution, denoted as ZCN^+^ (equation 2 in Methods). If the data set contains multiple cell types, we assume that the gene expression level across all the cells follows a mixture of ZCN^+^ distributions. The zero component is used to model the dropout event, and each gene has its own dropout rate (1 − *p_g_*). Besides random dropouts, a single cell experiment usually fails to capture low-expression genes if the sequencing depth is not enough. Thus, we use a multivariate Gaussian distribution censored below zero to model the “amplified” gene expression taking into account of the measurement limit. This Tobit model was also used in a previous study^18^ for the gene expression in scRNASeq. We did some model checking to show that the marginal distribution of most genes can be fitted by ZCN^+^ distribution (Supplementary Notes).

Usually the expressions of genes involved in a common biological function are correlated in a way such that the variability of gene expression can be summarized by the variation of several gene modules. A gene module can represent a pathway such that some genes in the pathway are co-expressed if the pathway is activated. Based on these intuitions, the co-variance matrix of gene expression for each cell type can be expressed as a low-rank structure plus idiosyncratic noises as in the factor analysis. The expression of a gene module in each cell is represented by a latent factor. These gene modules can be shared among closely related cell types. Thus, we reuse the same gene modules for each cell type but the activity of each gene module can be different to allow for gene modules either unique to a cell type or shared among several cell types. Since a gene module may only contain a few genes, we posit a Laplace prior for the loading matrix such that only some of the genes have nonzero weights for a gene module. This model enables us to utilize cell clusters and gene-gene correlation to impute the dropout values and further refine cell clusters and gene modules.

We could estimate the dropout rate for each gene according to the marginal distribution in scRNASeq. However, for low-expression genes, it is difficult to distinguish zero entries due to biological low-expression from dropout events. To avoid unduly imputation, we set an upper bound on the dropout rate *a priori* (see Methods). Nevertheless, with the help of bulk RNASeq data from similar cell population, we can estimate the dropout rate by combining the prior dropout rate and the ratio of the mean expression in the single cell data set with that in the bulk RNASeq. The overview of our model is shown in Figure 1. For inference, SIMPLEs employs a nested Monte Carlo EM algorithm and initializes the algorithm only using genes with fewer zero entries (see Methods). After estimating the model parameters, SIMPLEs runs several MCMC iterations and outputs the mean and variance of each imputed values as well as multiple copies of imputed expression matrix. The multiple imputations can be used to estimate the stability of clustering memberships.

**Figure 1.**
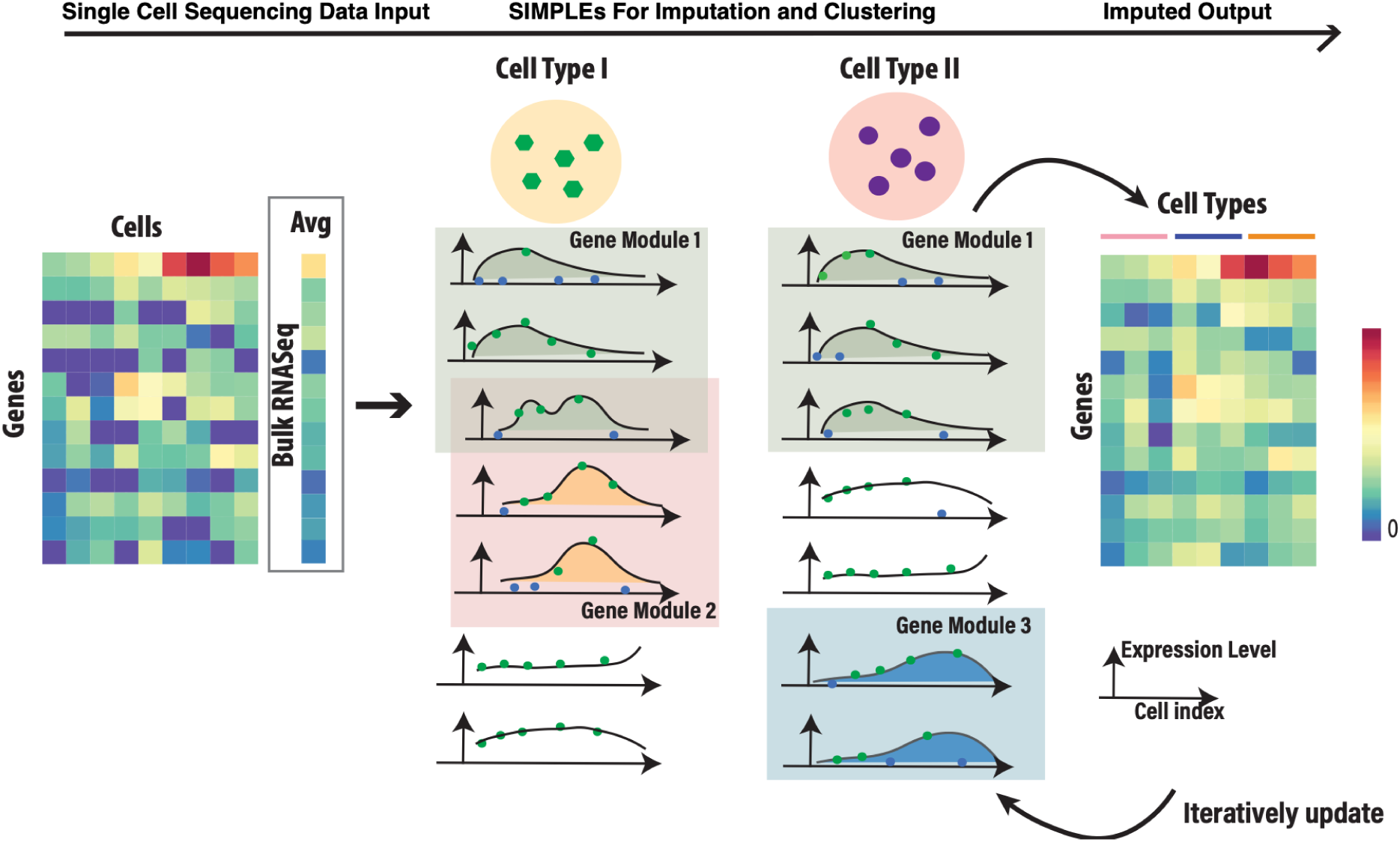
Overview of SIMPLEs. The input is a gene expression matrix from scRNASeq and, optionally, the corresponding bulk RNASeq data. SIMPLEs identifies the cell clusters and gene modules in which gene expressions are highly correlated. Each gene module is shown as a shaded box and the gene expression pattern across cells is represented by a curve. Each dot is the gene expression in a cell. Green dots are “amplified” expressions and blue dots are dropouts. Gene module 1 is activated in both cell types but others are unique to one cell type. The expression pattern of genes shared in modules 1 and 2 show characteristics of both modules. SIMPLEs outputs the imputed matrix, cell clusters and gene modules.

### Simulations

In the first experiment, we simulated data following the procedure in Li and Li^13^, in which the gene expression was sampled from mixtures of normal distributions and each gene was independent within each cluster of cells (see Methods). We simulated three cell clusters and randomly selected 20 cluster-specific genes for each cluster that are only differentially expressed in one of the cluster. Then, we randomly added dropouts to each gene by sampling from a Bernoulli distribution with gene specific dropout rate. We considered both high and low dropout rate scenarios in which dropout rates decay either slowly or quickly along with mean expression levels, respectively. The average dropout rate per gene is about 0.7 or 0.4 for the high and the low dropout rate scenario, respectively. The bulk RNASeq data were simulated as the mean expression of each gene.

We compared SIMPLEs with MAGIC, scImpute, VIPER, SAVER, and SCRABBLE, using the original data matrix without imputation as a baseline. To compare the performance, we measured the clustering performance after imputation using the adjusted rand index (aRI); the mean squared error (MSE) of imputed values compared with the truth; the ability of calling cluster-specific genes from the imputed data matrix using Area Under the ROC Curve (AUC). A larger aRI means that the clustering result is closer to the true labels, and aRI is 1 when the true labels are completely recovered. From Table 1, SIMPLE-B performed better than others; SCRABBLE and SIMPLE did the second best in all aforementioned performance evaluations. When the dropout rate was high, all methods except SCRABBLE and SIMPLEs were not able to identify the true cell types, resulting in poor aRI values. VIPER was not able to recover the true gene expression and its MSE of the imputed values was even worse than the original data. The imputed value of VIPER was often much smaller than the true value. VIPER also had the longest running time among all the methods.

**Table 1.** Performance of simulations with no gene correlation

|  | No impute | SIMPLE | SIMPLE-B | scImpute | MAGIC | SCRABBLE | VIPER | SAVER |
| --- | --- | --- | --- | --- | --- | --- | --- | --- |
| *Low dropout rate* |  |  |  |  |  |  |  |  |
| **aRI** | 0.88±0.03 | **1.00±0.00** | **1.00±0.00** | 0.93±0.02 | 0.85±0.05 | 0.99±0.00 | 0.96±0.02 | 0.95±0.02 |
| **MSE** | 2.52±0.02 | 0.86±0.02 | **0.78±0.02** | 1.04±0.01 | 1.08±0.01 | 0.81±0.01 | 1.87±0.03 | 1.71±0.02 |
| **AUC** | 0.91±0.01 | 0.95±0.01 | **0.96±0.01** | 0.93±0.01 | 0.59±0.01 | 0.94±0.01 | 0.90±0.01 | 0.92±0.01 |
| *High dropout rate* |  |  |  |  |  |  |  |  |
| **aRI** | 0.06±0.02 | 0.72±0.08 | **0.90±0.04** | 0.42±0.07 | 0.33±0.06 | 0.70±0.06 | 0.06±0.03 | 0.07±0.02 |
| **MSE** | 3.5±0.02 | 1.39±0.02 | **0.97±0.01** | 1.58±0.01 | 1.82±0.01 | 1.21±0.01 | 4.45±0.05 | 2.84±0.02 |
| **AUC** | 0.83±0.01 | 0.86±0.01 | **0.88±0.01** | 0.83±0.01 | 0.57±0.01 | 0.87±0.01 | 0.78±0.01 | 0.84±0.01 |

The numbers shown are mean±SD over 10 repetitions. Upper panel: low dropout rate; lower panel: high dropout rate.

SIMPLE-B, which incorporates the simulated bulk RNASeq information, was consistently better than SIMPLE, reflecting that the bulk RNASeq provides substantial information about the dropout rate that could not be reliably identified especially for lowly expressed genes. Figure 2 shows the projections of a few simulated low and high dropout data sets onto two-dimensional spaces, based on either the original data or the imputed data using different methods. SIMPLEs recovered the cell types even when cell clusters were indiscernible because of high dropout rate (Figs. 2f,h). Moreover, the performances of SIMPLEs were not sensitive to the number of factors chosen in the model (Supplementary Fig. 1). In this simulation, the true number of factors should be zero but the results shown were from a misspecified model where the number of factors was set at 2. The clustering performances of SIMPLEs were not sensitive to other parameters as well but the performances for imputation and identifying the true marker genes deteriorated if the prior upper bound for the dropout rate in SIMPLE was set much smaller than the true dropout rate (Supplementary Fig. 1). The impact of the dropout rate prior in SIMPLE-B was much lessened because of incorporating bulk RNASeq.

**Figure 2.**
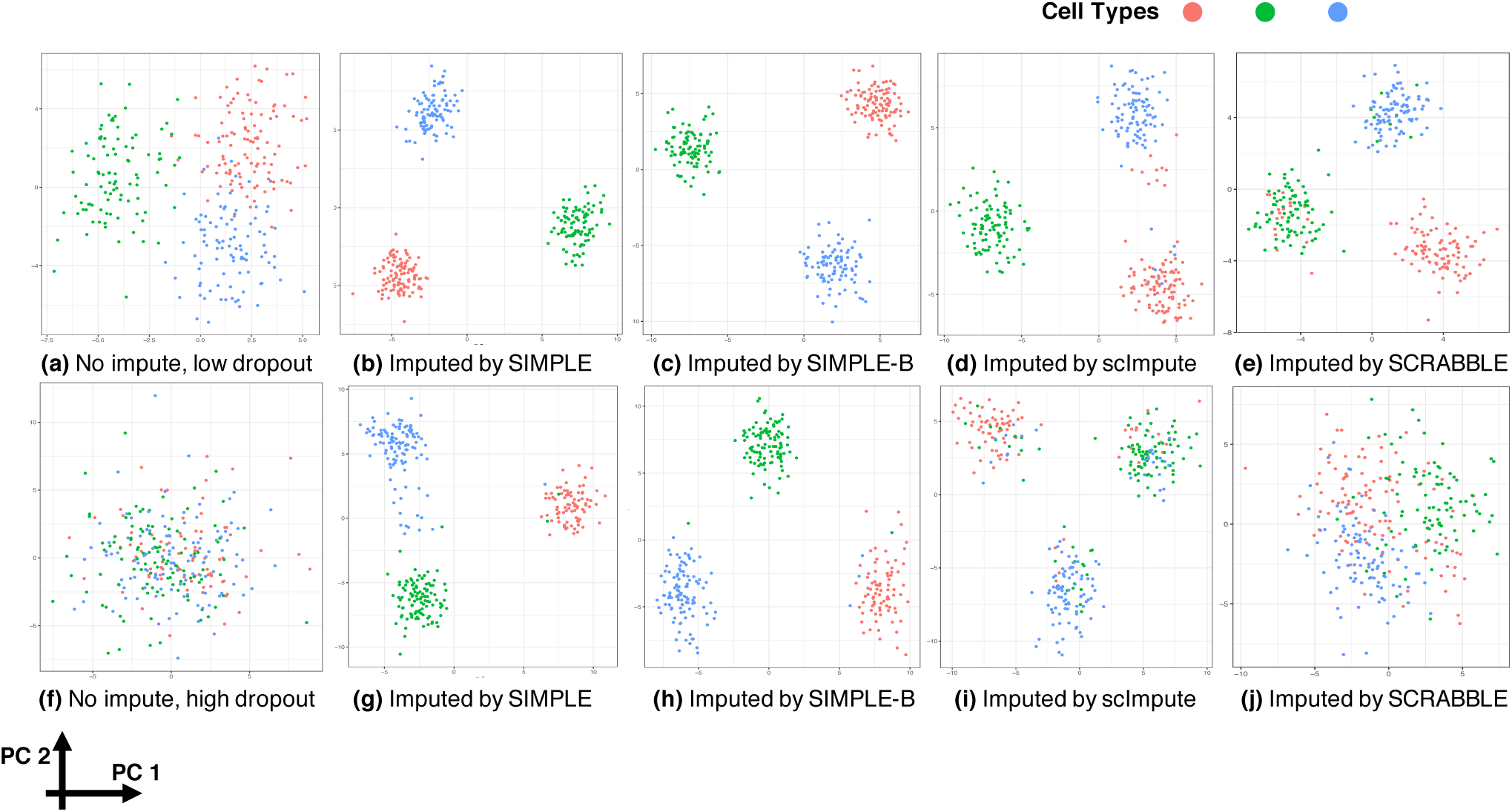
Examples of simulation data set. We projected both the original and the imputed data matrix onto the space spanned by the first two principal components (represented by the X and Y-axes, respectively). Each point is a cell colored by its true cell type color. (a)-(e): low dropout rate scenarios, and aRI = 0.85, 1, 1, 0.86, and 1 for respective methods. (f)-(j): high dropout rate scenarios, and aRI = 0.03, 0.81, 0.95, 0.30, and 0.80 for the respective methods. Other methods that were tested performed worse. (Table 1).

In the second experiment, we simulated data with correlated gene expression in each cluster of cells. The mean gene expression for each cluster was simulated in the same way as in the first experiment; the covariance matrix was the summation of two matrices: one for modeling the genes’ correlations within each module, and the other, a diagonal matrix, for modeling the idiosyncratic noise for each gene (see Methods). We considered two scenarios: (a) a large number of small sized gene modules with no overlapping genes; (b) a small number of large sized modules with some genes shared by a pair of modules. In addition, we simulated independently expressed genes so that the total number of genes was 1000, the same as in the previous simulation. Genes are positive correlated only if they are in the same module; otherwise, their correlation is zero. To focus on the effect of correlation between genes, the dropout rate was simulated the same as the low dropout rate scenario in the previous experiment (more details in Methods). Clustering and identifying marker genes are more difficult in the second scenario, as the within-cluster variations of gene expression are larger but the mean differences between clusters stay the same. However, large gene modules are beneficial for imputation using SIMPLEs since the method can incorporate correlated genes for imputing the missing entries. Besides the evaluation metrics proposed in the previous simulation, we also compared the gene-gene correlation estimated from the imputed data to the true correlation. Estimating correlations are of interest in some applications, such as constructing gene regulatory networks. To separate from the clustering performance, we only considered gene correlation within each simulated “true” cell cluster. The positive correlations between genes within a module and zero correlations in different modules were evaluated separately. Table 2 shows the MSEs between the estimated correlation from imputed data and the true positive or zero correlations, denoted as “cor1” and “cor0” respectively.

**Table 2.**
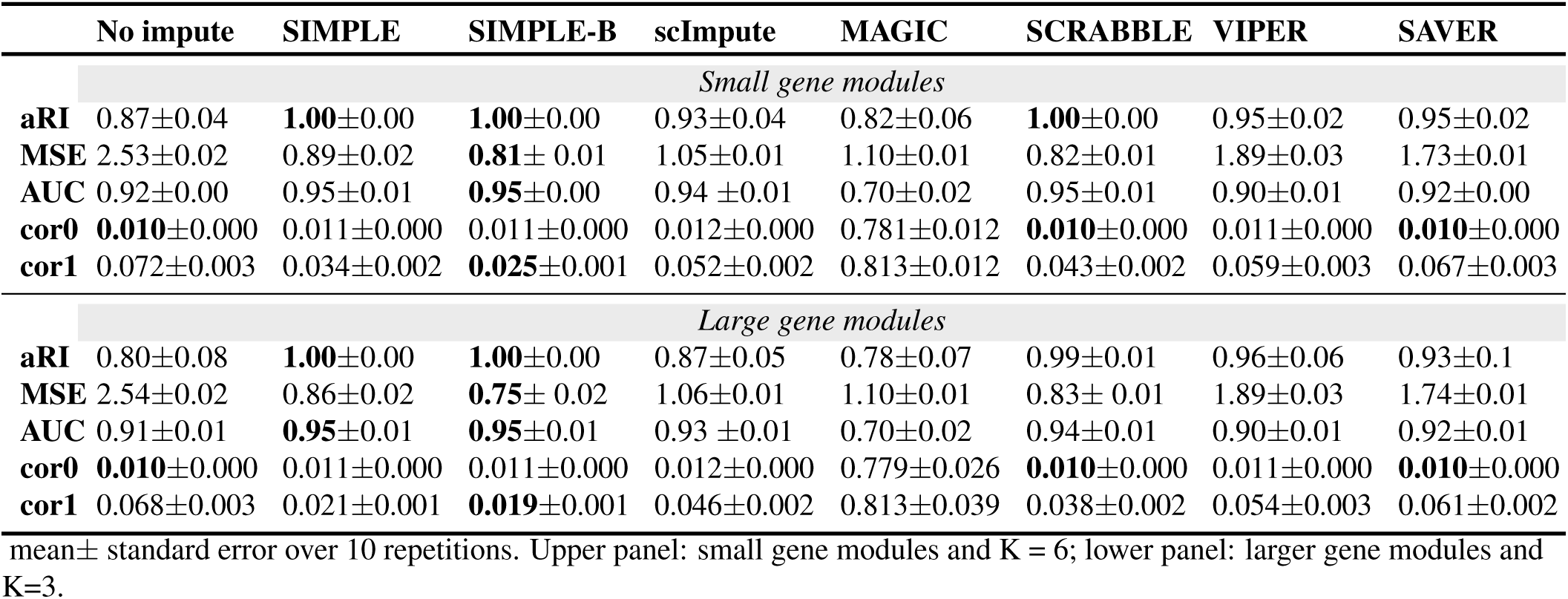
Performance of simulation with gene correlation

Most methods did well in clustering and identifying marker genes in this experiment since the dropout rate was relatively low (Table 2). MAGIC and scImpute were not as good as other methods in the large gene modules scenario because the strong correlations among the genes overwhelm the differences between clusters. SIMPLE-B had the smallest MSE of the imputed values and SCRABBLE performed the second best. Comparing large and small gene modules scenarios, SIMPLE-B and SIMPLE had smaller MSEs in the scenario with large gene modules, where more correlated genes can be used for imputation. Imputation MSEs of other methods were similar in these two scenarios. For estimating correlations within a cluster, SIMPLE was better than others even without incorporating bulk RNASeq. Because of random dropout, the sample gene-gene correlation matrix is usually smaller than the true one. SIMPLEs recovered the gene modules and restored the correlations between genes, but other methods often imputed gene expression close to the cluster mean and underestimated the non-zero within cluster gene correlations. MAGIC performed much worse than other methods. Due to its forcing the imputed gene expressions to follow a common trend, MAGIC substantially overestimated gene correlations.As a consequence, MAGIC also performed poorly in cell clustering and identifying differentially expressed genes.

We varied the tuning parameters of SIMPLEs for this experiment and observed similar results as in the first experiment (Supplementary Fig. 2). All of the performances were not sensitive to the number of factors (*K*). The performance was reported using the true *K* in Table 2, but stayed almost the same for other Ks. We provided some guidelines for choosing *K* in practice in the Methods section.

### Human embryonic stem cell differentiation

We applied SIMPLEs to a study of human embryonic stem cell (hESC) differentiation towards definitive endoderm^19^. It includes a single cell RNASeq dataset for seven cell types (1018 cells in total), referred to as “hESC cell types”: two types of embryonic stem cells (H1 and H9), definitive endoderm cells (DEC), endothelial cells (EC), human foreskin fibroblasts (HFF), neuronal progenitor cells (NPC) and trophoblast-like cells (TB), and the bulk RNASeq dataset for each cell type. DEC, EC, and NPC are differentiated cells from three germ layers. DEC and EC share a transient precursor state called mesendoderm. In this study, they also conducted another scRNASeq experiment at different time points in the differentiation process towards DEC, referred to as “hESC time course”, and produced the corresponding bulk RNASeq data at each time point.

First, we compared the clustering performance of SIMPLEs with other methods for the “hESC cell types” data set. We only input the average gene expression across all cell types in the bulk RNASeq to SIMPLE-B and SCRABBLE. Otherwise, the gene expression from bulk RNASeq can disclose the true cell type information we wanted to recover. However, for real application, we can definitely incorporate the bulk RNASeq of different cell types for better imputation and identification of differential expressed genes among different cell types. This data set has good quality with low dropout rate. The cells can be nearly perfectly clustered using either the original data without imputation or imputed ones by SIMPLEs, VIPER, or SAVER. However, the aRI was only about 0.75 using scImpute, MAGIC, and SCRABBLE (Fig. 3b). Since SAVER was too conservative to impute the data (also shown in the following experiments), it is not surprising that it had similar clustering performance as using the original data without imputation.

**Figure 3.**
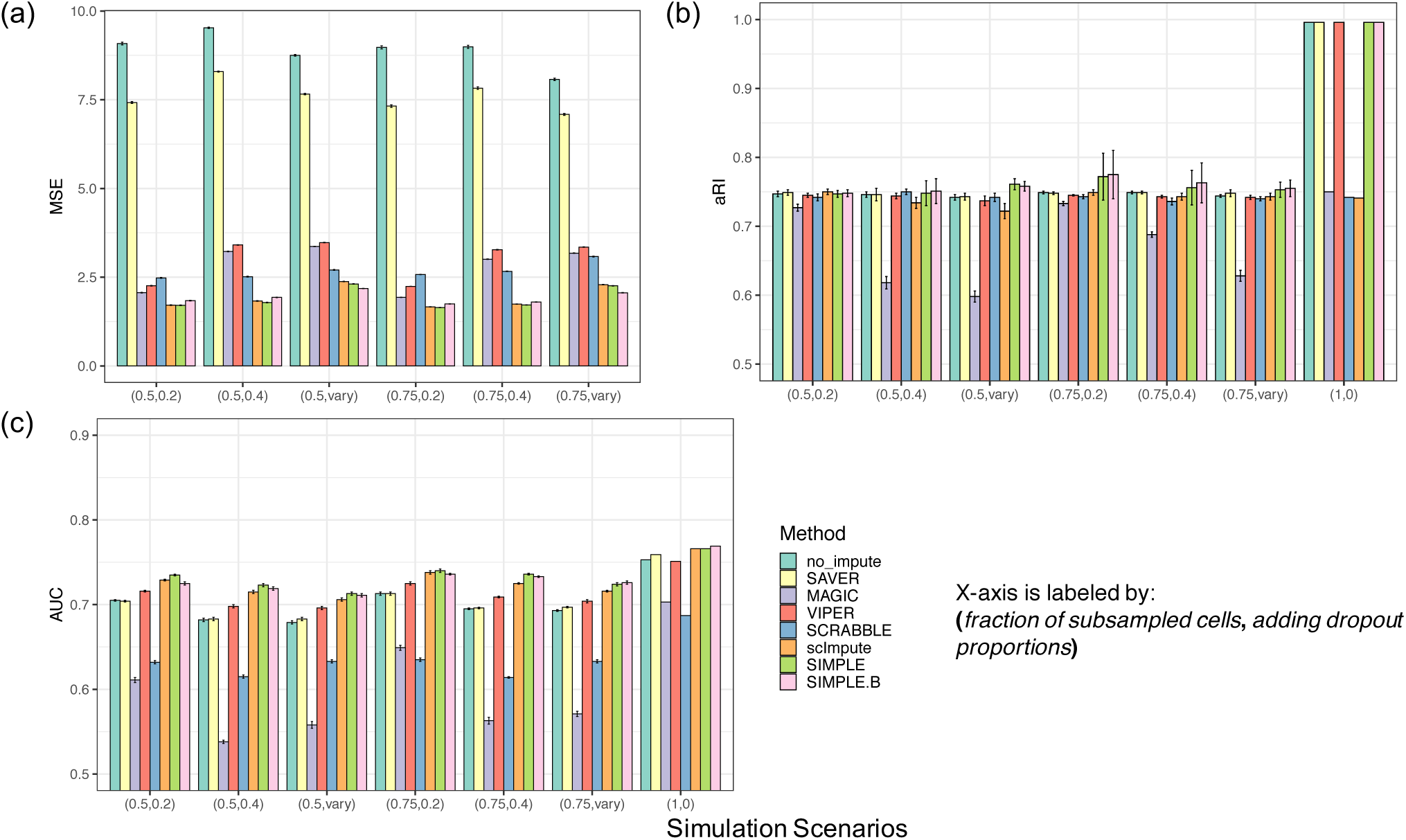
Performance comparisons for various simulation scenarios based on the hESC cell types data. (A) mean squared error of imputed values compared with the original values; (B) aRI comparing Kmeans clustering results with the true cell labels; (C) AUC comparing differentially expressed genes identified by different imputation methods with the genes identified by bulk RNASeq using DESeq2^20^as ground truth (see Methods). Each bar represents the result from a imputation method or data without imputation (“no_impute”); X-axis marks different simulation scenarios. The first number in the parenthesis is the fraction of cells subsampled and the second is either the dropout probability adding to the original data set or “vary” for cases where the dropout rate per gene decreases with mean expression. The error bar is the standard error over 10 repetitions.

A main utility of scRNASeq data in comparison with bulk RNASeq is to explore cell heterogeneity within major cell types. From the distributions of the latent factors (F) in each cell type, we observed several factors showing larger variations in DECs than in other cell types (Supplementary Fig. 5a), indicating hyper transcriptional stochasticity especially for genes in the corresponding gene modules. As an illustration, we re-performed imputation and clustering for DECs and ECs only, based on top 1000 genes with largest absolute weights in gene module 1. We showed the results from SIMPLE without involving additional information from bulk RNASeq to distinguish DEC and EC. DECs and ECs were still well separated but both of them can be further divided into subtypes (Supplementary Fig. 5b). The expressions of these top genes further validate the distinct transcriptomes of cell subtypes (Supplementary Fig. 5d). The heatmaps using the original data and the imputed ones look similar since the dropout rate is low for this data set, yet the differences among subtypes were more discernible after imputation (Supplementary Fig. 5c-d). We observed similar subtypes and gene modules by SIMPLE-B.

Then, as another evaluation of the imputation methods in consideration, we took the original hESC cell types data set as the ground truth and added dropouts to evaluate how well each method can recover the original expression matrix. We designed two dropout schemes: 1) randomly select 20% or 40% entries and set them zero, uniformly for

2 all genes; 2) set the probability of dropout decreasing with the mean expression of each gene, i.e., 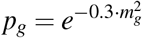. We also subsampled 50% or 75% of all the cells and 3000 genes uniformly in order to test how the performance is influenced by the sample size and to examine the stability of the performance as data vary due to subsampling. For this experiment, SIMPLEs performed the best in almost all scenarios for all the evaluation metrics (Figure 3). scImpute performed the second best and its clustering results were about same as that of SIMPLEs. The imputed data by SAVER incurred almost the same mean squared error as the original unimputed data, as SAVER was too conservative to impute any dropouts (Figure 3a). None of the methods could get perfect clustering result (Figure 3b). As we subsampled genes and cells for this experiment, the variance of aRI (adjusted rand index) was large for some cases. To test the performances of identifying differentially expressed genes, we used the marker genes obtained from bulk RNASeq as the ground truth. Although the AUC was higher for identifying marker genes using SIMPLEs, the markers identified from single cell data sets and bulk RNASeq only agreed to some extent, as the AUC was around 0.75 when we used the original data set without additional dropouts (Figure 3c). Comparing with SIMPLE, SIMPLE-B showed better performance incorporating bulk RNASeq when we added artificial dropouts whose rate is a monotonically decreasing function of the mean expression (Figure 3a). Furthermore, we varied the parameters in SIMPLEs (Supplementary Figs. 3-4). The clustering performance and identifying differential expressed genes were not sensitive to the choices of parameters. The imputation performance was worse if specifying too small the upper bound of the dropout rate for SIMPLE, but the prior dropout rate did not affect the imputation performance for SIMPLE-B. Moreover, when assuming a large number of factors and a small penalty parameter in the prior of the loading matrix, the model was susceptible to over-fitting, as the imputation mean squared error was larger.

Finally, in order to show that the imputed expression levels by using SIMPLEs can reconstruct gene expression trends in biological processes, we also applied SIMPLEs to the hESC time course data set. It contains 758 single cells captured at 0, 12, 24, 36, 72 and 96h in the cell developmental process from pluripotent state through mesendoderm to DE, as well as bulk RNASeq data at each time point. Since cells’ developmental process is asynchronous, cells collected at the same time could be at different developmental stages. Thus, the true developmental state of each cell should be correlated but not exactly the same as the cell’s time stamp. Clustering results using either the original data or the imputed data indicated that similar cell types are observed at 72h and 96h, whereas cells from other time points are well separated (Supplementary Fig. 6). Cells from 72h and 96h form two clusters, of which one contains purely cells at 96h, but the other is composed of the cells from both time points, indicating that cells entered the final stage of differentiation asynchronously. As a consequence, in order to identify gene markers for each developmental stage, it is not sufficient to compare gene expression for cells at each time point from bulk RNASeq. On the other hand, single cell data provides information of the developmental stage of each cell and can be used to identify key genes governing the developmental process.

If the imputation reflects the true biological process, the expression level of the known marker genes in the developmental process should be correlated well with the developmental stage. Considering the fact that the true developmental stage of each cell is not known, we applied Monocle^18^ to order the cells using the original data. It assigned a pseudo-time to each cell indicting the developmental stage of the cell. The pseudo time agreed with the true time label from 0h to 36h. However, cells from 72h and 96h cannot be distinguished by pseudo-time. Then, we checked the imputed gene expression of several known markers along the pseudo-time (Fig. 4, Supplementary Fig. 7). Imputed values by SIMPLEs followed the cell developmental process and preserved the variability of gene expressions in a single cell, while other methods (e.g., scImpute and MAGIC) tended to impute the gene expression as the mean expression in each cell cluster. Although SIMPLEs utilizes correlated expression changes of the genes with similar functions to infer the dropout values, the imputation will not interfere genes with different functions that can be turned on or off at different stages. For example, *PRDM1*, a DE-specific gene, was expressed at a high level after 72h indicating that cells differentiated toward the DE state, whereas pluripotent state marker *NANOG* was down-regulated during differentiation^19^. MAGIC changed all the expression values to the mean in each cell cluster and completely ignored the heterogeneity of single cell expression. SCRABBLE mis-identified the cell state and imputed *PRDM1* by the overall mean expression before 12h when *PRDM1* is not expressed (Fig. 4a). Imputed values by VIPER were correlated with the cell differentiation timeline for most of the cells, but had a similar weakness as SCRABBLE in that it imputed the expression of *PRDM1* for the cells at the very early stage as high as the ones at the intermediate stage. Compared to VIPER, SIMPLEs imputed zero entries at a relatively low expression level for *PRDM1* in the cells at early stages. Moreover, SIMPLEs preserved the stochasticity of single cell expression while other methods reduced the variability of gene expression after imputation. SIMPLEs estimates the variance of expression for each gene. For genes with large variances, the probability of observing zeros from the amplified component is high, so SIMPLEs imputes less frequently and retains low expression level for zero entries. For example, *NANOG*, a pluripotent state marker, was expressed at a relatively low level and had a high variance in cells at late stages. VIPER imputed the expression of *NANOG* by its mean after 36h, but SIMPLEs imputed zero entries at low level and maintained the variability of gene expression at late stages (Fig. 4b). More imputed expressions of the marker genes along with the cell developmental pseudo-time are shown in Supplementary Fig. 7, demonstrating that SIMPLEs can faithfully recover the gene expression pattern designated by the gene’s biological functions.

**Figure 4.**
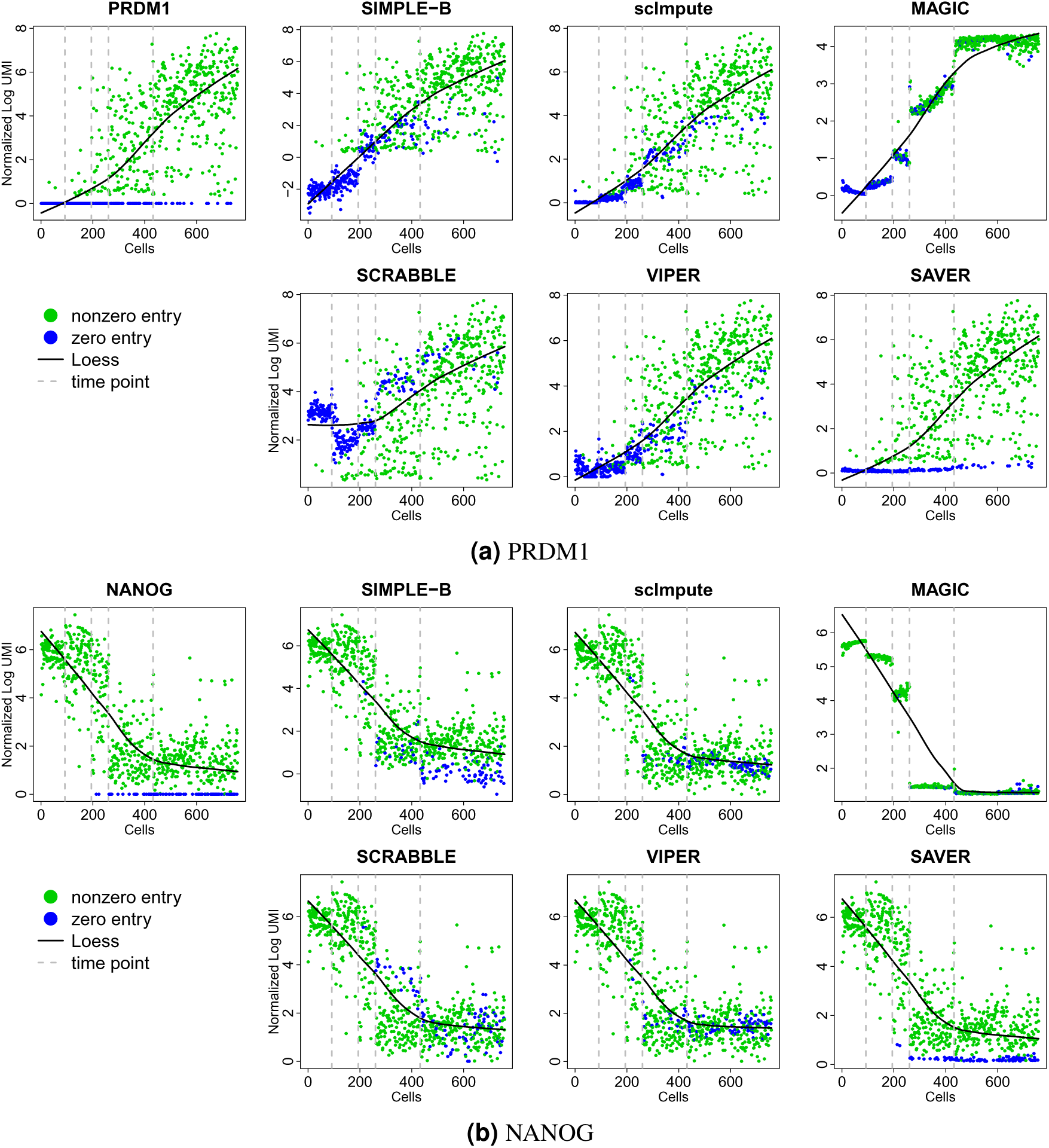
Examples of imputed marker gene expressions along the hESC developmental pseudo-time. Each dot is the gene expression level in a cell. Green dot: nonzero expression in the original data. Blue dot: zero in the original data but is imputed by different methods. The solid black lines are the smoothed loess curves computed for each data. Cells are first ordered by time label, then within each time stamp, are ordered by pseudo-time output by Monocle, except that we treated cells at 72h and 96h as one group and ordered them by pseudo time. The dashed lines are the boundaries of 0h, 12h, 24h, 36h cells.

Moreover, SIMPLEs can restore the (anti-)correlation between genes. For example, both *CER1* and *GATA4* are early DE-specific markers, which were turned on as early as 36h in the DE differentiation process. However, the correlation between them was attenuated because of dropout. Nevertheless, the close relationship between these two genes was revealed after imputation by SIMPLE-B (Fig. 5a). For MAGIC, most pairs of genes had extremely high correlations after imputation. scImpute and VIPER also did well, but imputation from SCRABBLE and SAVER underestimated the correlation between *CER1* and *GATA4*. As another example, *T* and *CXCR4* are negatively correlated as cells transform from a T-positive to a CXCR4-positive state during differentiation towards DE^19^. The negative correlation between *T* and *CXCR4* was weakened in the original data due to dropouts. All of the imputation methods retrieved the negative correlation but to different extent, from the strongest correlation by MAGIC, then by SIMPLE-B, to the weakest by SAVER which is the same as the original data (Supplementary Fig. 8). Furthermore, we identified differentially expressed genes at each time point based on the imputed data (see Methods). From the gene expression heatmap, cells at different stages showed distinct transcriptome profiles (Fig. 5b). These differentially expressed genes of each developmental stage are putative key regulators controlling the developmental process. They included known regulators of the developmental process, such as pluripotent state markers: *SOX2, POU5F1, NANOG, DNMT3B*; early cell state markers (expressed in 12-24h of differentiation): *NODAL, ID1, T, MSX2*; late cell state markers (turned on at 36-72h of differentiation): *CER1, DKK4*; and DE markers: KIT, *PRDM1, POU2AF1*.

**Figure 5.**
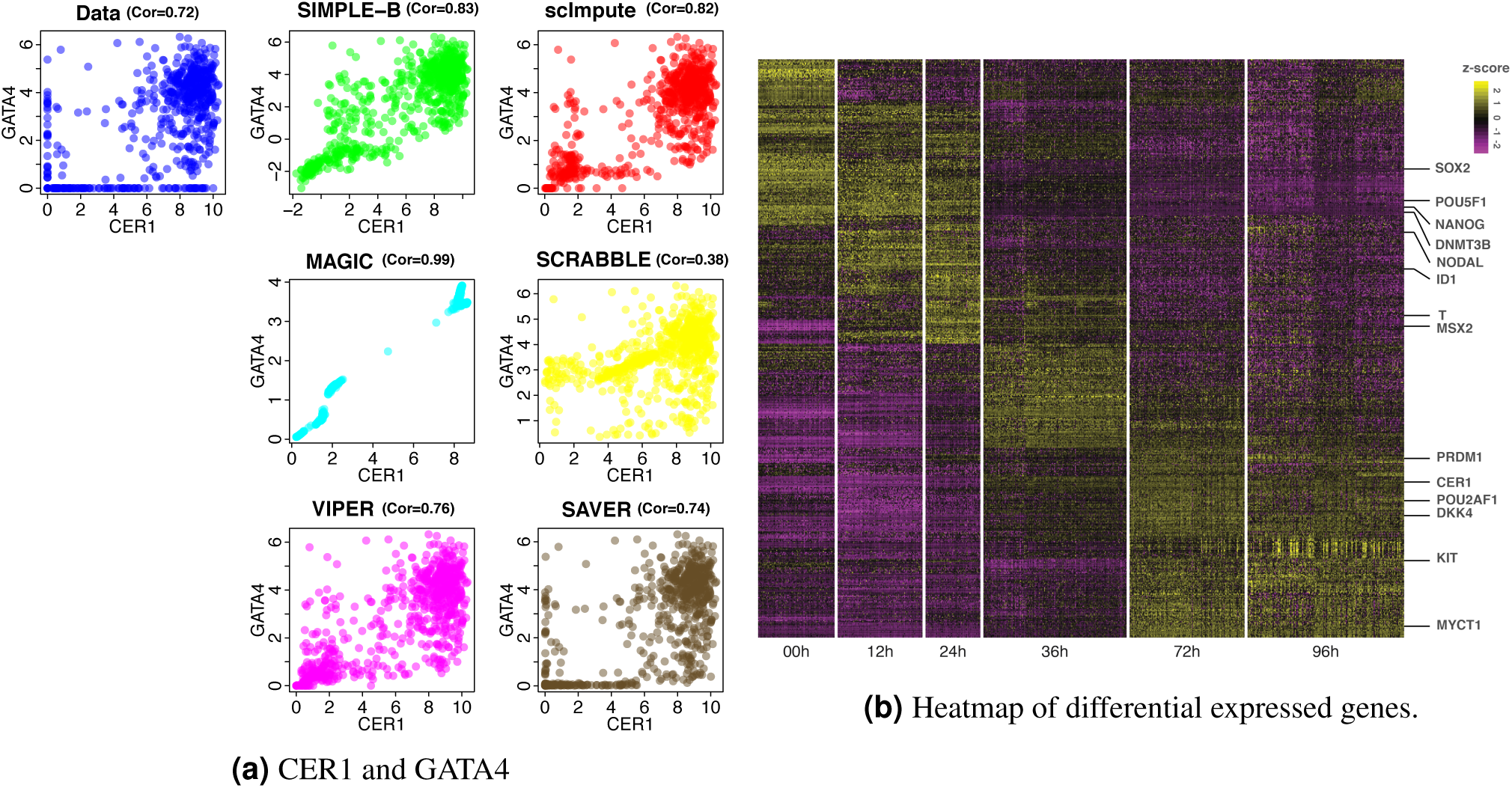
hESC time course data. (a) Original or imputed expression of CER1 and GATA4 by different methods. Each dot represents a cell and the correlations between the two genes are shown in the title of each sub-figure. (b) Imputed gene expression by SIMPLE-B. Each row is a gene and each column is a cell ordered by the time label. The color indicates the z-score (centered and scaled by gene). The heatmap shows top 100 differentially expressed genes ranked the p-values from Wilcoxon rank sum test for each time point.

### Mouse preimplantation embryos

As another example, we applied our method to a single cell data set of mouse embryos^21^. It contains 12 zygotes, 22 cells at the 2-cell stage, 14 at 4-cell, 36 at 8-cell, 50 at 16-cell and 133 blastocysts (267 cells in total). The data set does not have the corresponding bulk RNASeq, so we only compared SIMPLE with other methods. Using either the original or the imputed data by various methods, we derived similar cell clustering results, which separated blastocysts into two clusters but merged 16-cells and 8-cells into one cluster, indicating that blastocysts is even more diverse than cells at 8-cells and 16 cells stages (Fig. 6). Indeed, these major cell stages can be further divided into 10 subtypes^21^. Although different imputation methods had similar results for clustering the major cell stages, SIMPLE can distinguish subtypes better than others (Fig. 6). When trying to cluster the cells into 10 subtypes, SIMPLE achieved aRI=0.8, whereas the second best method SAVER had aRI = 0.6. In contrast, the clustering aRI was only 0.4 without imputation. Some subtypes have very few cells, e.g. less than 10, rendering them unrecognizable by any method. Yet this data set contains many more cells at the blastocyst stage, which can be further divided into three subtypes. SIMPLE could clearly separate three stages of blastocysts, i.e., early, mid blast, and late blastocyst, while other methods missed the substructure of cells because the imputation might have over-smoothed the gene expression. In addition, SIMPLE can output multiple imputations and compute the stability of the clustering result for each cell. For example, a small group of 8-cells separated from others on the t-SNE plot have a high clustering uncertainty as they sometimes were clustered as an isolated group while sometimes joined other 8-cells; clusters of 8-cells and 16-cells were also not stable as they were often clustered together but occasionally separated into two clusters (Fig. 6).

**Figure 6.**
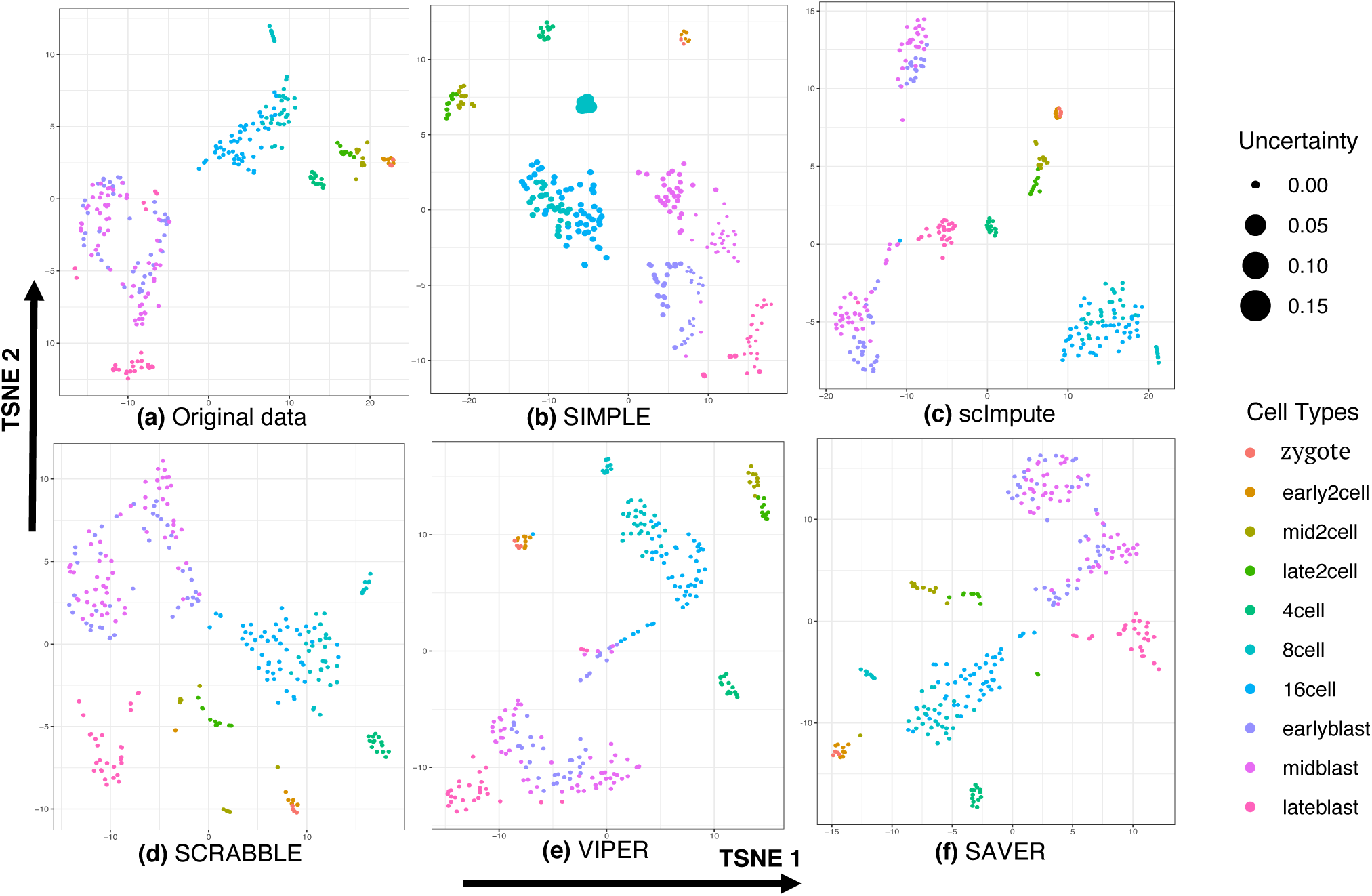
Visualizing the original data or imputed data by different methods using t-SNE for the mouse embryos data set. Each point is a cell colored by subtypes. For SIMPLE, the size of the point indicates the uncertainty of clustering membership of each cell across multiple imputations (the larger the size, the more uncertain the cell is). For SIMPLE, we varied the number of factors from 5 to 8 and the results were similar. The result shown here was obtained from *K* = 8. For SIMPLE and scImpute, we set the number of clusters equal to 6, which is the number of major cell stages.

Looking into the distributions of cells’ latent factors in each subtype, we observed that subtypes of cells can be discriminated by several latent factors (Supplementary Fig. 10). For example, factors 2 and 3 can separate late and early blastocysts from the rest of blastocysts, which explains why SIMPLE was able to distinguish different stages of blastocysts. Factor 6 can distinguish different stages of 2-cell; and factor 1 can differentiate 8-cell and 16-cell. Other factors did not show significant differences in any particular subtype, indicating that the expression of associated gene modules vary uniformly across all subtypes. As shown above, SIMPLEs models within cluster covariance structure and can discover different subtypes beyond major cell types.

Then, we compared the expressions of marker genes for each subtype before and after imputations. To identify marker genes, we merged some of the subtypes with less than 10 cells in the original study, and obtained 8 subtypes: zygote, 2-cell, 4-cell, 8-cell, 16-cell stage, and early-, mid-, late-blastocyst. Marker genes were identified by comparing their expression levels in each subtype with the rest of the cells using the imputed data (see Methods). The imputed expression of marker genes by SIMPLE showed clearer patterns of specifically expressed in one or more subtypes than that of other methods (Fig. 7; imputed gene expressions by other methods are shown in Supplementary Fig. 9).

**Figure 7.**
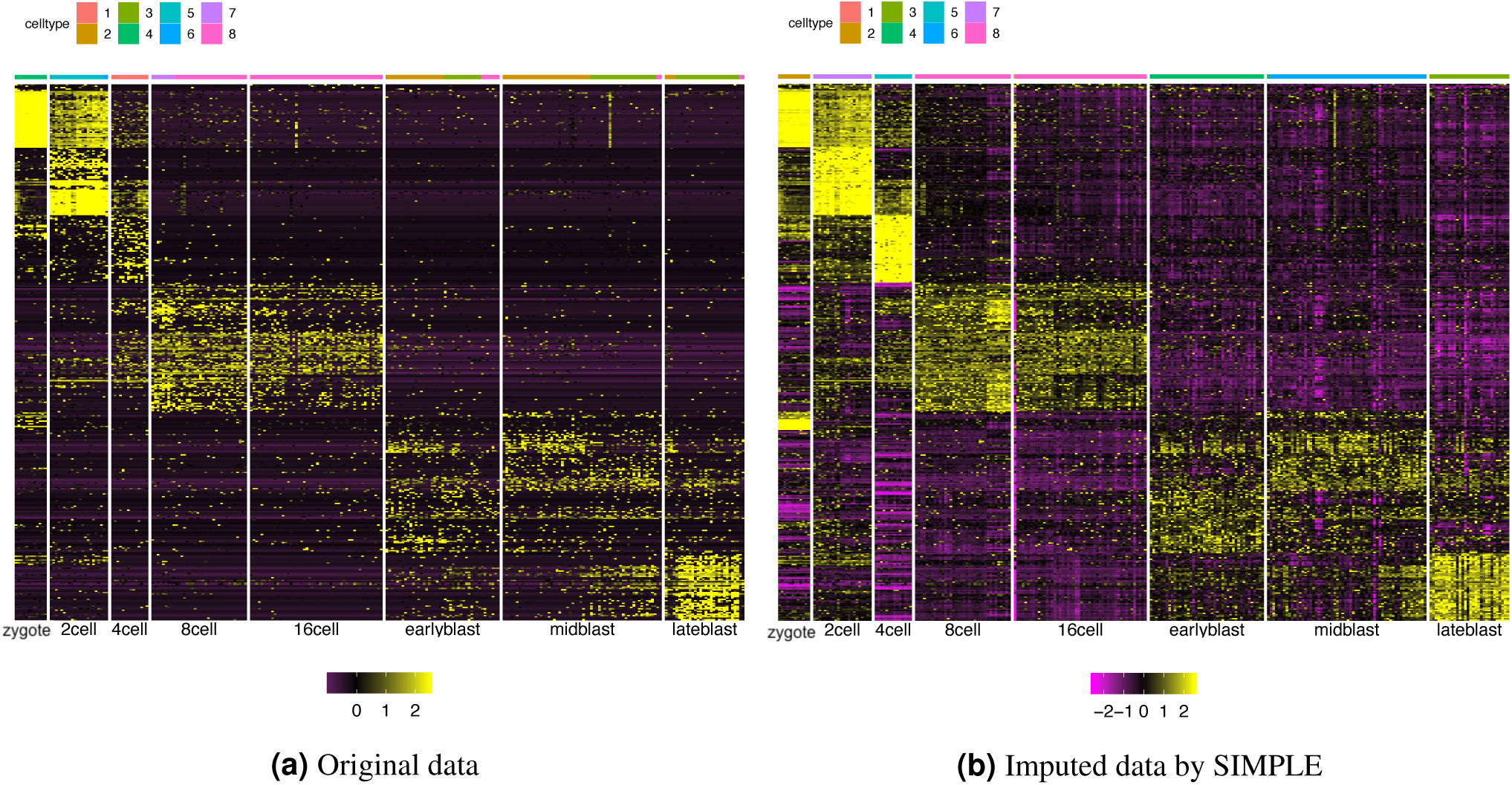
Gene expression values based on the mouse embryos data set (A) without imputations, and (B) with imputations produced by SIMPLE. Each row is a marker gene and each column is a cell ordered cell types. The purple to yellow color indicates the z-score range (centered and scaled by gene). The color bar above the heatmap shows the clustering result by each gene expression matrix. The order of the genes is the same in these heatmaps, but the cells within each cell type are ordered by the clusters obtained using each data set. Differentially expressed genes for each of the 8 cell types were identified by Wilcoxon rank sum test using imputed data. For clarity, we show the top 50 marker genes for each cell type ranked by the p-values.

### Mouse immune cells from multiple organs

To show the scalability of SIMPLE on large number of cells and more diverse cell types, we applied SIMPLE on 12,905 immune cells from 12 mouse organs^22^ including 22 known immune cell types. The preprocessing procedure is described in Methods section. The original data and the data with imputations obtained by SIMPLE were visualized using t-SNE in Figure 8, where colors indicate different cell types. Results from imputed data resembled the cell clusters identified from the original data. The overlapping of cells with different labels, especially using the original data, is mainly due to similar cell types in the same lineage on the cell ontology. These cell types come from different tissues but has the same ancestor on the cell hierarchy (Supplementary Figs. 11 and 12). Compared with the original data, the imputed data showed tighter clusters for cells from the same type, but separated different cell types further apart, e.g., the immature B cell from marrow and B cell mainly from other tissues.

**Figure 8.**
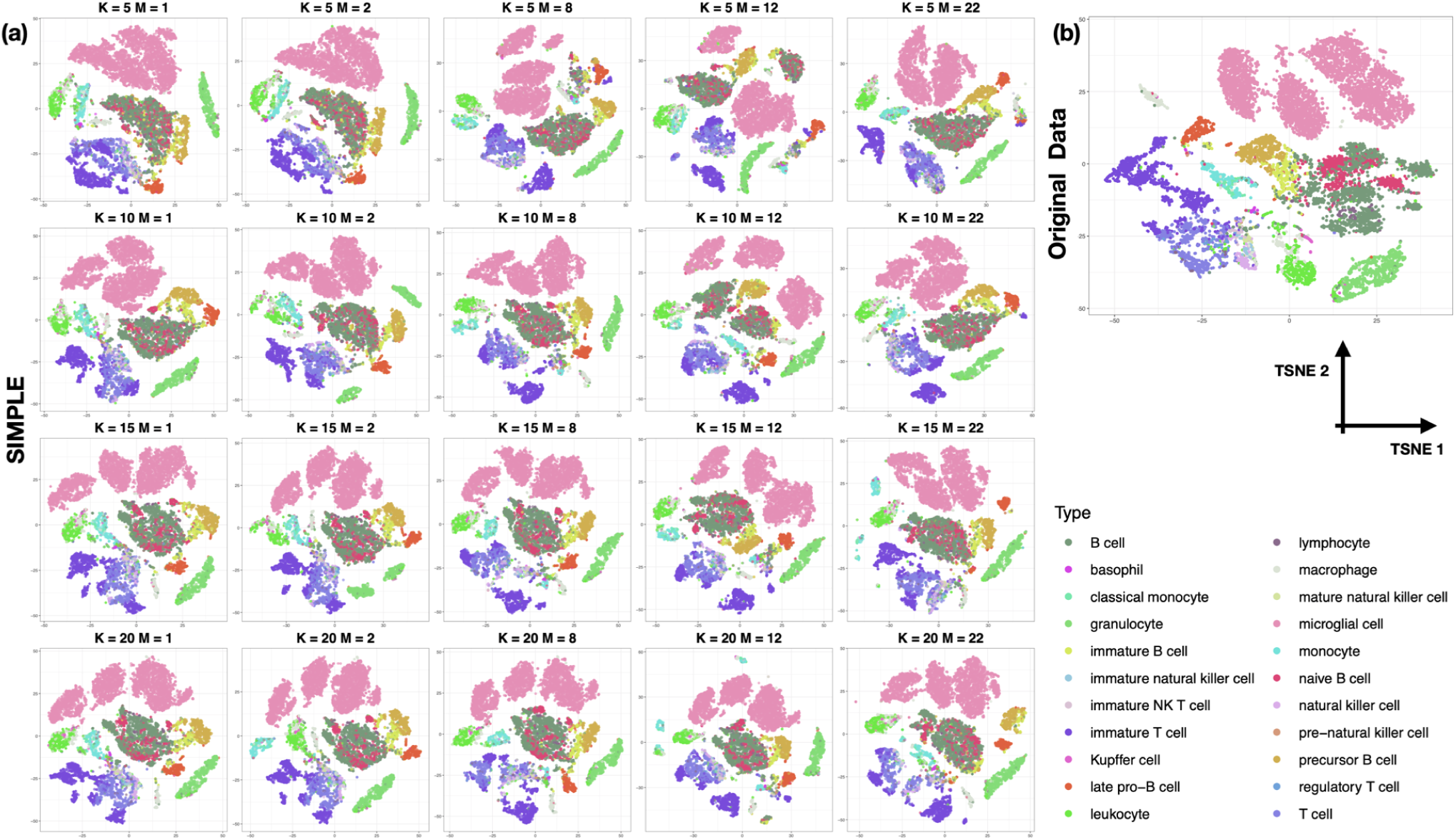
t-SNE based data visualization of both the original data and the imputed data by SIMPLE with different parameters. Each dot represents a cell and colors indicate different cell types. (a) The visualization of imputed data by SIMPLE using different combinations of *K* = 5,10,15,20 and *M* = 1,2,8,12,22. (b) The visualization of the original data.

Varying the parameters in SIMPLE, i.e., the number of clusters (M) and the number of factors (K), did not change the t-SNE plot using the imputed data much, reflecting the robustness of SIMPLE with respect to parameter choices. Based on these observations, we suggest to set *M* as 1 and *K* as 10 for imputation and preliminary exploration of a data set, which is the default setting of SIMPLEs without prior knowledge of the consisting cell types. Nevertheless, increasing either *K* or *M* can reveal subtypes of the cells. As shown in Figure 8, when increasing *M*, different types of T cells reflecting the organ of origin becomes more discernible: immature T cells mostly from thymus and marrows are separated T cells from other organs such as fat, lung, limb, muscle, and spleen (labeled as T cell) and immature NK T cells.

Furthermore, we analyzed the latent factors and their associated genes modules captured by SIMPLE for all types of T cells when *M* = 1 and *K* = 10. We identified representative genes for each gene module based on the weights in the corresponding column of the loading matrix *B* (see Methods) and the biological functions of these genes. Distributions of several latent factors (e.g. medians) showed significant differences across six organs (Supplementary Fig. 13), indicating that these gene modules identified by SIMPLE can reflect the origin of T cells. We took two of the gene modules corresponding to latent factors 3 and 9 as examples, since both of them had large average weights over genes and the latent factors showed distinct patterns of distributions across organs (Figure 9a). The medians of latent factor 9 varied in most of the organs considered, indicating differentially expression of gene module 9 in various organs; on the other hand, the medians of latent factor 3 in cells from thymus was higher than that of other tissues, suggesting specific expression of gene module 3 in thymus. These distinct patterns imply that the genes in these two gene modules play different roles in T cell developments.

**Figure 9.**
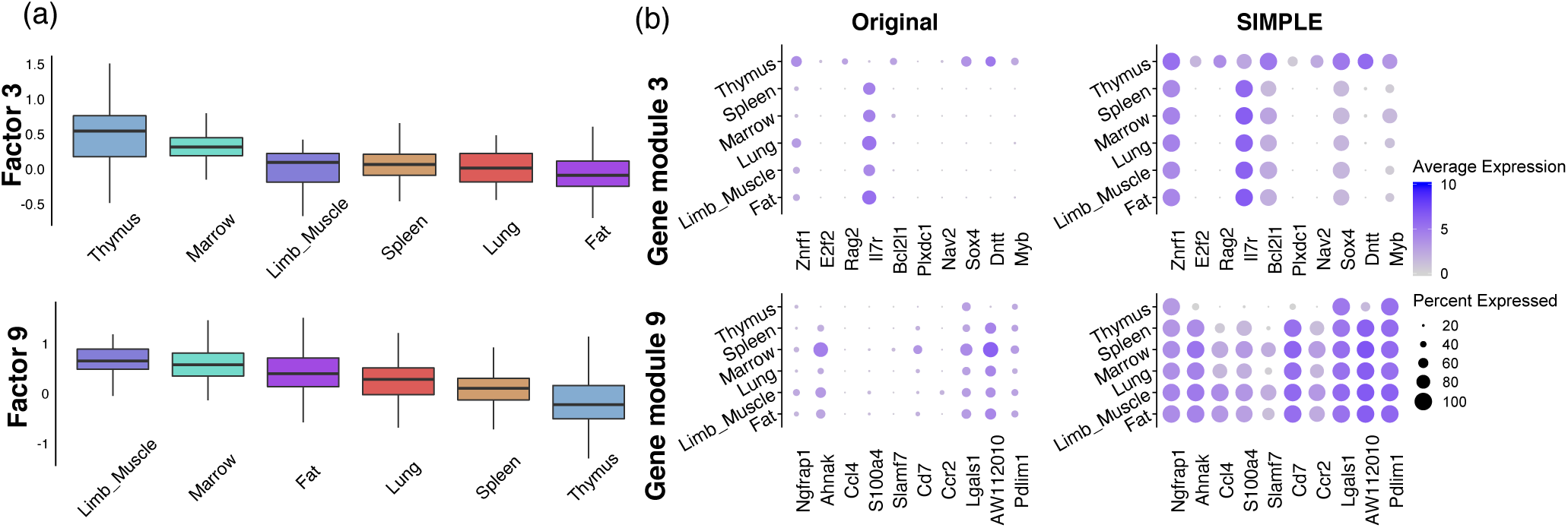
Analysis of the latent factors 3 and 9 discovered by SIMPLE for T cells in six organs. (a) The boxplots show the distributions of factors 3 and 9 in six organs. (b) The dot-plots show the gene expression patterns of the top 10 genes across six organs in the original and imputed data. These genes have largest coefficients in the corresponding gene module.

To verify the above observation, we showed top 10 genes with the largest loadings in these two modules (Figure 9b). After imputation, the percentages of cells expressing these top ranked genes increased as expected, yet the imputed data maintained gene expression variation across organs. Among the top genes from the gene module 9, *S100a4* showed higher expression in fat and limb muscle, which is consistent with the result from the previous study^22^. In the meanwhile, many of the top genes from gene module 3 had higher expression levels in thymus and function in the early stage of T cell development. For example, *Myb* encodes c-Myb transcription factor, and is essential in early T cell development^23^; *Dnnt* encodes a DNA nucleotidylexotransferase, which is a specific DNA polymerase in pre-B and pre-T cells; finally, *Rag2* involves in the recombination during B and T cell development.

## Discussion

SIMPLEs impute dropout values in the single cell RNASeq data based on both cell similarities and gene correlations. The imputed data matrix can be used to reduce dimensionality for visualizing the cell spectrum, to identify markers between different samples, and to construct gene co-expression networks. SIMPLEs iteratively cluster the cells, identify correlated gene modules and impute dropout values within each cluster utilizing the expressions of other correlated genes. Integrating with the corresponding bulk RNASeq data, SIMPLE-B can give a better estimate of dropout rates, which influences strongly how much the data should be imputed. It is shown in simulations that SIMPLE-B combining bulk RNASeq has significant advantages over SIMPLE in recovering the dropout values (e.g. Table 1). Only one very recent method, SCRABBLE, can incorporate bulk RNASeq information for imputation. Compared with SIMPLE-B, SCRABBLE is less optimal in estimating gene-gene correlation and in cell clustering when high dropout rates are present. SCRABBLE also underestimates the variability of gene expressions in single cell, as shown in our analysis of the hESC data set.

Most existing imputation methods only consider single imputation, whereas SIMPLEs can output multiple imputations, which can be used to assess clustering stability and reliability (Fig. 6). To the best of our knowledge, only SAVER can output the variance of each imputed value, but it does not reflect the joint variability of multiple genes’ expressions. Furthermore, SIMPLEs can identify activated gene modules with large variability in one or more cell types, which can be used to identify subtypes of cells or gene modules that are associated with the attributes of the samples. Our analyses of the first two real data sets illustrate that latent gene modules discovered by SIMPLEs can be used to further classify subtypes of human definitive endoderm cells and stages of blastocyst cells in mouse embryos respectively. Moreover, in the human embryonic stem cell data, we showed that our imputed values for marker genes followed the developmental stages of each cell and can be used to discover candidate regulators that are important to the developmental process. Finally, the analysis of a large-scale dataset on mouse immune cells from multiple organs demonstrates the scalability and robustness of SIMPLEs. We also identified several gene modules that varied in expression in T cells from multiple organs, including key genes that regulate T cell development.

Since the likelihood function of our model is non-convex, the solution of the algorithm depends on the initial clustering and imputation. Thus, we only selected high quality genes to estimate the cell clusters and latent factors during initialization. As shown by Jin et al.^24^, in high dimensional settings when the number of features is much more than the number of samples, feature selection is crucial for recovering the true clusters. To further improve clustering performance, more sophisticated gene selection procedure may be adopted for initialization^25^. We used a nested EM algorithm for estimation, but optimizing the factor loading matrix *B* in the M-step was computational intensive. In order to analyze large-scale single-cell sequencing data, a stochastic gradient descent algorithm can be employed to reduce the computational time in optimizing *B*.

We provided some guidelines in choosing parameters for SIMPLEs and also showed that SIMPLEs are not sensitive to the number of factors, and the priors of loading matrix and dropout rate. In the mouse immune cells data set, we showed that the imputation results were robust to the input number of clusters, so we set *M =* 1 by default for imputation and visualization if no cell type information is available. Determining the number of clusters in the data is a challenging task. Since the number of cells in each cell type and the distance between cell types vary, the number of clusters chosen by some criteria, such as maximizing the likelihood, is often different from biological knowledge. Moreover, the number of clusters chosen for imputation should also depend on the dropout rate. If the dropout rate is high, imputation requires pooling information from more cells so the number of clusters should be small and vice versa. After initial imputation and visualization, one might rerun SIMPLEs with different *M*s for clustering or apply other clustering algorithm and visualization method on the imputed gene expression matrix. We only implemented a simple clustering method as we focused on imputation in this paper, but more sophisticated clustering method may be necessary to improve the clustering accuracy for rare cell types^26^ and cell types with a complex hierarchy.

In our present model, we did not incorporate the uncertainty of estimating *B*, which is relevant for quantifying the uncertainty of the imputed values. *B* can be estimated accurately when the number of cells increases, but a more careful study of the uncertainty in estimating *B* with the present of large amount of missing data is needed. Moreover, SIMPLE-B relies on the fact that the cell composition in the bulk RNASeq is the same as in the single cell experiment, which might be violated in some real data. One possible approach is to characterize cell type compositions in bulk RNAseq data by deconvolution with gene expression of each cell type in the single cell data based on high quality genes^27^. Finally, the relationship among genes and cells can be much more complicated than the linear latent factors model assumed in SIMPLEs. Since neural networks have shown promises in modeling complex nonlinear relationships and general probability distributions^28^, it is of interest to incorporate neural networks into our model to capture certain nonlinear aspects of the data, which may enhance our understanding about the interactions of cell clusters and gene functional modules at the single cell level.

## Methods

### Details of the model and inference procedure

The log-normalized gene expressions in each single cell are modeled by a zero-inflated negative-censored multivariate normal distribution. Suppose there are *M* cell types, for cell *i* of type *C_m_*, its vector of gene expressions, 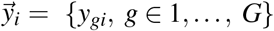, is modeled as:

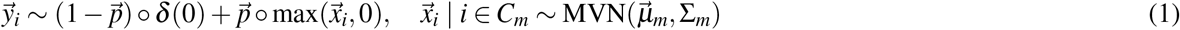

where 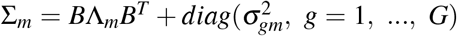 with {Λ_*m*_ ≡ diag(*λ_km_,k* = 1,…,*K*), *m* = 1,…,*M*} denoting a sequence of *K* × *K* diagonal matrices satisfying 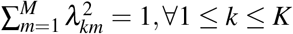; notation “○“ denotes the entry-wise product; and “MVN” stands for the multivariate Normal distribution. Let 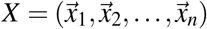 be the gene expression matrix without truncation and dropout. The purpose of imputation is to recover *X* from *Y*. The first component of Eqn (1) is a point mass at zero, which models the dropout event, and 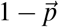 is the dropout rate vector, where 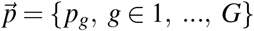. Each gene has its own dropout rate and we assume that the dropout rate does not depend on the cell type. The second component, called “amplified”, models the measurement limit of low-expression genes in the single cell experiment. It follows a multivariate normal censored below zero, denoted by 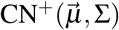, is defined as: 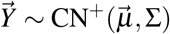, if 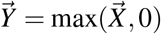, where the maximum is taken for each entry of 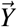, and 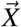 follows multivariate normal distribution with mean 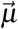 and covariance Σ. Using this notation, equation 1 can be rewritten as:

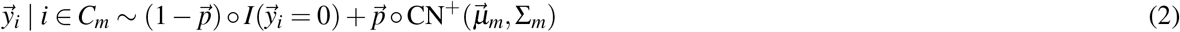

We call the distribution in Eqn (2) a zero-inflated negative-censored multivariate Gaussian distribution (ZCN^+^). 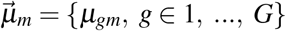 is the vector of average expression of every gene in cell type *m*. When *μ_gm_* is smaller, we expect more zero counts generated from the second component. For real applications, we set a threshold *δ*_0_ close to zero and regard the expression below *δ*_0_ as “zero”, allowing for some background noise (typically *δ*_0_ = 0.1). The co-variance matrix (∑_*m*_) of gene expression for each cell type can be expressed as a low-rank structure (*B*Λ*_m_B^T^*) plus idiosyncratic noises (*diag*(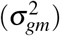)). The *loading matrix B* = [*B_gk_*] is a *G* × *K* matrix where *K* is the number of gene modules or pathways. We used the same *B* for all the clusters allowing gene modules shared in different cell types, but the activity of each gene module may vary. Without noise, each gene expression can be represented by a linear combination of these gene modules, where each row of *B* is the vector of weights for each gene to be mapped to the gene modules. The activity of the gene modules in a cell type is controlled by Λ_*m*_. When gene module or pathway *k* is unique to a single cell type, e.g. *m*_0_, *λ_km_, m* = 1,…,*M* is non-zero only for cell type *m*_0_. Closely related cell types usually have similar gene modules’ activities. Since the log-likelihood does not change when both *B* and Λ_*m*_ are scaled by reciprocal amount, we constrain the sum of 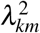 for each factor in different cell types to be 1 (Eq. 4).

To facilitate imputation and clustering, we augment the data by *Z*, the indicator of cluster membership, and *F*, the latent factors for each cell:

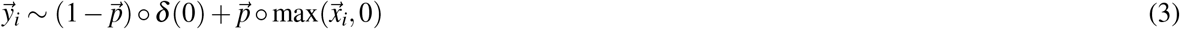

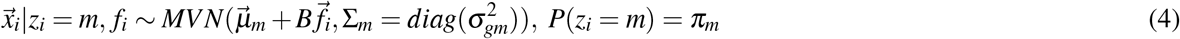

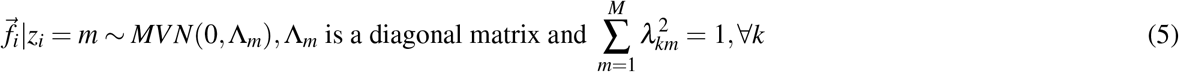

The factors are arranged as a *K* × *N* matrix, *F*, where row 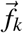. is the factor associated with gene module *B_·k_*. Each column 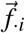 of *F* represents the expression of the gene modules in cell *i*, which is i.i.d given the clustering membership. We use a Laplace prior for *B* so that the weight of each gene within a gene module is sparse. Non-zero entries in each column of *B* reflect the correlation between genes imposed by a gene module. We rank genes representing a gene module by their (non-zero) weights in the corresponding column of loading matrix *B*. In the application below, we assume that each gene has the same “noise” variance for each cluster, i.e. the 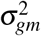 are the same for all *m*.

The EM algorithm is often used to estimate parameters in missing data problems and for models with a latent structure. However, the E-step of the EM algorithm needs to compute the expectation of the log-likelihood over all the latent variables, which cannot be achieved directly for our model. Thus, we employed the nesting Monte Carlo EM algorithm^29^, which alternatively uses Gibbs sampling to impute missing data from its conditional distribution at current parameters, computes the expectation over the indicators (*Z*) and factors (*F*) conditioning on the imputed data (*X*), and updates the parameters (*B*, Λ, Σ) by maximizing the approximated expectation of full log-likelihood.

The algorithm is initialized with *B* = 0, implying that each gene is independent of others for cells in the same cluster. For genes with many zero entries, it is hard to distinguish dropout from amplified component based only on the marginal distribution of individual gene expression. Thus, we only use genes with a small number of zero entries (“high-quality” genes) for initial imputation. Then, we estimate factors *F* using “high quality” genes, and project the expression of the remaining genes onto *F*. Here we define “high-quality” genes as those whose percentages of zeros are lower than a threshold and also set a minimal number of genes for initialization. We found that initializing *F* using only high-quality genes is better than using all the genes, since information about the major structure of cells, i.e. the cell cluster and latent factors, can be revealed from only a subset of genes. Initial imputations for genes with many zeros can be quite noisy and including these genes can drive the estimation towards undesirable local modes.

If without bulk RNASeq, we fit each gene expression by a zero-inflated negative-censored Gaussian distribution and obtain the maximum likelihood estimator of the dropout rate (1 − *p_g_*). We set an upper bound on dropout rate to prevent excessive imputation, since we do not have enough information to estimate dropout rates for “low quality” genes. With bulk RNASeq, we can estimate the dropout rates more accurately. We assume that bulk RNASeq measures the mean expression level for each gene without dropout, then the dropout rate per gene is estimated by a linear combination of the prior dropout rate and the ratio of mean expression of scRNASeq and that of bulk RNASeq if the scRNASeq and the bulk RNASeq are normalized similarly. Including a prior dropout rate is to alleviate the instability of the ratio when the gene expression value in the bulk RNASeq is too low. We used 1 − *p_m_* to denote the upper bound or the prior dropout rate, which can be estimated experimentally^30^ or empirically from the data set (see Implementation details). To account for possibly different scales, we assume that a gene’s expression in bulk RNASeq is a linear transformation of its mean expression in the scRNASeq. Since highly expressed genes are unlikely to be dropped out, we conducted a weighted least squares method to estimate the scaling factor with weights proportional to the square of the mean expression value. If the input data have cell type labels, we estimate the dropout rate for each cell type separately. For more details, see Supplementary notes.

### Clustering and identifying marker genes after imputation

Since the imputation methods we compared with do not output the clustering result directly, we simply used Kmeans after projecting the imputed or the original data onto the space spanned by the top *k* principal components, e.g., *k*=20, of the data matrix. Since no imputation methods provide an explicit procedure to select the number of clusters, we used the true number of clusters for Kmeans. For SIMPLEs, we set M to be the number of major cell types or the number of time points in hECS time course data set, and used the cluster memberships directly output by SIMPLEs in a majority of experiments, except for clustering subtypes of cells. For clustering DEC and EC cells in the hESC cell types data set, we set *M* = 7 for imputation, and then did Kmeans clustering on the imputed data for DEC and EC cells; for the mouse embryos data set, we set *M* = 6 for imputation, and then did Kmeans clustering on the imputed data for identifying subtypes. If we have multiple imputed expression matrices (with nearly independent imputations), we apply the aforementioned clustering procedure for each imputed expression matrix and record the frequency that each pair of cell is in the same cluster, forming a co-clustering consensus matrix^31^ based on which we assign an uncertainty score for each cell (Supplementary Note).

To identify marker genes, we applied Student’s t-test for each gene in the first two experiments and the Wilcoxon rank sum test for the real data sets, comparing the imputed gene expression in one cell cluster with the rest. To separate from the clustering error, we used the true labels of the cells for identifying cluster-specific genes. Then, we tested the genes with fold change > 1.2 and ranked the genes by the p-values output from t-test or Wilcoxon test. To compute the AUC, we compared the test statistics to the truly differentially expressed genes for each cluster, and obtained the average AUC. For the first two experiments, we know the true markers for each cluster. For the hESC cell type data set, we identified differentially expressed genes for each of the seven cell types using DESeq2^20^ from bulk RNASeq and treated the genes with FDR < 1*e* − 6 and fold change > 1.4 as the true markers. The number of cell type specific markers varied from 2000 to 4000 for the seven cell types.

### Simulation procedures

For the first two experiments, we set the number of genes *G* = 1000 and the total number of cells *N* = 300 divided into three clusters evenly. We simulated the mean expression of each gene by *μ_g_* ~ log – Normal(0.5,0.5). Then, we randomly selected 20 cluster-specific genes for each cluster, and changed the mean expression in one of the three clusters. The fold change was sampled uniformly from {0.2,0.5,1.2,1.5,2} to make the overall mean expression of cluster-specific genes similar to other genes. After that, we sampled the gene expression in each cell following the multivariate normal distribution with a specified mean and covariance matrix. Finally, we added dropout to each gene by sampling from the Bernoulli distribution with dropout rate 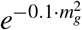 or 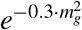, corresponding to high or low dropout rate scenarios, where *m_g_* is the overall mean expression for each gene. The mean dropout rate of every gene was about 0.7 or 0.4 for high and low dropout rate scenario respectively. We simulated bulk RNASeq by taking the mean expression of each gene, i.e. *m_g_*.

For simulating independent gene expression within cluster, we sampled the gene expression in cell cluster *m* i.i.d. from 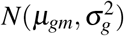, where the standard deviation *σ_g_* ~ Gamma(2,0.3). To simulate correlated gene expression, we constructed the covariance matrix as a low rank matrix plus a diagonal matrix, i.e. Σ = *BB^T^* + Σ_0_, where Σ_0_ is a diagonal matrix with each diagonal entry sampled from Gamma(2,0.3), *B* is a *G* × *K* matrix, *K* is the number of factors (or gene modules), and *BB^T^* induces a overlapping block structure reflecting co-expression of gene modules. We considered two scenarios: a large number of small gene modules, where we set K=6 with 32 genes in each module with no genes shared by two modules; a small number of large gene modules with more genes in multiple modules, where we set K=3 with 100 genes in each module. In the second case, 55 genes are shared by two consecutive modules given an order of the modules. Each entry of *B* is either 1/4 or 0. In addition, we simulated about 800 independently expressed genes so that we had *G* = 1000 in total. The correlation between genes is stronger in the second than that in the first scenario. The norm ‖*BB^T^*‖_*F*_ is larger in the second case and the largest eigenvalue of *BB^T^* is 11.4 compared to 2 in the first case. For simulation with correlated gene expression, we only considered the low dropout rate scenario.

For the simulation of additional dropouts based on the hESC cell type data set, we either randomly set 20% or 40% entries to zero uniformly for all genes or set the probability of dropout decreasing with the mean expression of each gene, e.g., 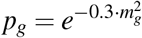. We also subsampled 50% or 75% of all the cells, which correspond to 509 and 763 cells, respectively, and randomly selected 3000 genes that were expressed in more than 50 cells for each experiment to reduce the computational cost and test the stability.

### Implementation details

For the first two experiments, the results shown in the main text were obtained by setting the upper bound or the prior of dropout rate to be 1 − *p_m_* = 0.6 in SIMPLEs and scImpute. For simulations based on the hESC cell type data, the results shown in the main text were obtained by setting 1 − *p_m_* = 0.3 when we added 20% more dropouts and 1 − *p_m_* = 0.5 for other scenarios. For real applications, we set 1 − *p_m_* = 0.2 as default. The number of factors was set at 10 unless otherwise stated. The parameter in the Laplace prior of the loading matrix (*γ*) was set to be 1 unless otherwise stated. These parameters are typical choices for SIMPLEs and we did not tune these parameters further. Results varying different parameters in the simulations are shown in Supplementary Figs. 1-4. We set *δ*_0_ = 0.1 for SIMPLEs in all the experiments, and input the true number of clusters (*M*) to both SIMPLEs and scImpute except for the mouse immune cells data set where we varied *M*.

For real data applications, we recommended choosing genes with their fraction of zero entries less than 50% but keeping a minimum of 2000 genes for initialization. For the prior of dropout rate, we adopted the empirical Bayes approach, which first estimates the dropout rate per gene based on the marginal distribution for SIMPLE and bulk RNAseq for SIMPLE-B, then set 1 – *p_m_* as the median of the dropout rate across all the genes. For choosing the penalization parameter (*γ*) for weights in the loading matrix and the number of factors (*K*), we suggest to add some dropouts to the data set and find the optimal parameters to recover the original entries as we did in the simulations based on the hESC cell type data set. Typically, the penalization parameter should be small when the number of factors is small. Also, if the dropout rate is high, the imputation requires gene modules to be relatively large, such that more correlated genes can be incorporated for imputation.

For VIPER, SCRABBLE and SAVER, we used the default parameters for all the data sets. We used Monocle 2 to order the cells in the hESC time course data set and obtained pseudo time using the DDRTree algorithm.

### Data sets and softwares

We downloaded human embryonic stem cell differentiation and mouse preimplantation embryos data sets from https://hemberg-lab.github.io/scRNA.seq.datasets. For the hESC data sets, we selected genes with nonzero entries in at least 10% cells. After log normalization and further filtering, we included 8148 genes with the standard deviation greater than 1.2 in the hESC cell type data set and 5135 genes with the standard deviation greater than 1.5 in the hESC time course data set. Then, for the mouse embryos data set, we filtered out genes with percentage of zero entries less than 5%, which are more likely housekeeping genes that expressed in most of the cells, or greater than 60%, and used the remaining 8648 genes for further analysis. Finally, the mouse immune cells data set was obtained from a collection of single-cell RNASeq experiments of more than 100,000 cells from 20 mouse organs and tissues^22^. We extracted all the immune cells from FACS-based scRNASeq data for our analysis. The FACS scRNASeq data was downloaded from https://s3.amazonaws.com/czbiohub-tabula-muris/TM_facs_mat.rds. The cell annotation was obtained from the original study: https://raw.githubusercontent.com/czbiohub/tabulamuris-vignettes/master/data/TM_facs_metadata.csv, which contains the cell ontology class for each cell. Based on it, we collected cells with cell ontology ID CL:0000738 or its descendants from the cell hierarchy. The cell ontology data was downloaded from https://raw.githubusercontent.com/obophenotype/cell-ontology/master/cl-basic.obo. According to the data preprocessing procedure in the original paper, we excluded cells with fewer than 500 detected genes or larger than 20,000 detected genes, or cells with fewer than 50,000 reads or larger than 2 × 10^6^ reads. Moreover, we filtered genes in the immune cells if they are only expressed in less than 5% cells or the standard deviations of the log-normalized values are less than 1.2. Finally, 7,809 genes and 12,905 cells are kept in our analysis. These cells are from 12 different tissues, and can be classified into 22 immune cell types. SIMPLEs is available at https://github.com/JunLiuLab/SIMPLEs.

## Acknowledgements

We thank Professor Junying Yuan for inspiring discussions. This research is supported in part by NSF DMS-161303 and NIH 1RF1AG055521-01A1.

## Author contributions statement

Z.H. designed the method SIMPLEs. Z.H. and S.Z. conceived and conducted the experiments, and analyzed the results. J.L. supervised the study. All authors have reviewed the manuscript.

## Additional information

*Conflict of Interest: none declared*.

## references

1. Papalexi, E. & Satija, R. Single-cell RNA sequencing to explore immune cell heterogeneity. Nat. Rev. Immunol. 18, 35 (2017).

2. Villani, A.-c. et al. Supplementary Materials for. Sci. (80-.). 16, DOI: 10.1126/science.aah4573 (2017).

3. Zheng, G. X. et al. Massively parallel digital transcriptional profiling of single cells. Nat. Commun. 8, 1–12, DOI: 10.1038/ncomms14049 (2017).

4. Zeisel, A. et al. Cell types in the mouse cortex and hippocampus revealed by single-cell RNA-seq. Sci. (80-.). 347, 1138–1142, DOI: 10.1126/science.aaa1934 (2015).

5. Macosko, E. Z. et al. Highly parallel genome-wide expression profiling of individual cells using nanoliter droplets. Cell 161, 1202–1214, DOI: 10.1016/j.cell.2015.05.002 (2015).

6. Poulin, J. F., Tasic, B., Hjerling-Leffler, J., Trimarchi, J. M. & Awatramani, R. Disentangling neural cell diversity using single-cell transcriptomics. Nat. Neurosci. 19, 1131–1141, DOI: 10.1038/nn.4366 (2016).

7. Keren-Shaul, H. et al. A Unique Microglia Type Associated with Restricting Development of Alzheimer’s Disease. Cell 169, 1276–1290.e17, DOI: 10.1016/j.cell.2017.05.018 (2017).

8. Trapnell, C. et al. The dynamics and regulators of cell fate decisions are revealed by pseudotemporal ordering of single cells. Nat. Biotechnol. 32, 381–386, DOI: 10.1038/nbt.2859 (2014).

9. Klein, A. M. et al. Droplet barcoding for single-cell transcriptomics applied to embryonic stem cells. Cell 161, 1187–1201, DOI: 10.1016/j.cell.2015.04.044 (2015).

10. Shalek, A. K. et al. Single-cell RNA-seq reveals dynamic paracrine control of cellular variation. Nature 510, 363–369, DOI: 10.1038/nature13437 (2014).

11. Stegle, O., Teichmann, S. A. & Marioni, J. C. Computational and analytical challenges in single-cell transcriptomics. Nat. Rev. Genet. 16, 133 EP – (2015). Review Article.

12. van Dijk, D. et al. Recovering Gene Interactions from Single-Cell Data Using Data Diffusion. Cell 174, 716–729.E27, DOI: 10.1016/j.cell.2018.05.061 (2018).

13. Li, W. V. & Li, J. J. An accurate and robust imputation method scImpute for single-cell RNA-seq data. Nat. Commun. 9, DOI: 10.1038/s41467-018-03405-7 (2018).

14. Chen, M. & Zhou, X. VIPER: variability-preserving imputation for accurate gene expression recovery in single-cell RNA sequencing studies. Genome Biol. 19, 196, DOI: 10.1186/s13059-018-1575-1 (2018).

15. Peng, T., Zhu, Q., Yin, P. & Tan, K. SCRABBLE: single-cell RNA-seq imputation constrained by bulk RNA-seq data. Genome Biol. 20, 88, DOI: 10.1186/s13059-019-1681-8 (2019).

16. Huang, M. et al. SAVER: gene expression recovery for single-cell RNA sequencing. Nat. Methods 15, 539–542, DOI: 10.1038/s41592-018-0033-z (2018).

17. Miao, Z., Deng, K., Wang, X. & Zhang, X. DEsingle for detecting three types of differential expression in single-cell RNA-seq data. Bioinformatics 34, 3223–3224, DOI: 10.1093/bioinformatics/bty332 (2018). http://oup.prod.sis.lan/bioinformatics/article-pdf/34/18/3223/25731617/bty332.pdf.

18. Trapnell, C. et al. The dynamics and regulators of cell fate decisions are revealed by pseudotemporal ordering of single cells. Nat. Biotechnol. 32, 381–386, DOI: 10.1038/nbt.2859 (2014).

19. Chu, L. F. et al. Single-cell RNA-seq reveals novel regulators of human embryonic stem cell differentiation to definitive endoderm. Genome Biol. 17, 1–20, DOI: 10.1186/s13059-016-1033-x (2016).

20. Love, M. I., Huber, W. & Anders, S. Moderated estimation of fold change and dispersion for RNA-seq data with DESeq2. Genome Biol. DOI: 10.1186/s13059-014-0550-8 (2014).

21. Deng, Q., Ramsköld, D., Reinius, B. & Sandberg, R. Single-cell rna-seq reveals dynamic, random monoallelic gene expression in mammalian cells. Science 343, 193–196, DOI: 10.1126/science.1245316 (2014). https://science.sciencemag.org/content/343/6167/193.full.pdf.

22. Schaum, N. et al. Single-cell transcriptomics of 20 mouse organs creates a Tabula Muris. Nature 562, 367–372, DOI: 10.1038/s41586-018-0590-4 (2018).

23. Allen, R. D., Bender, T. P. & Siu, G. c-myb is essential for early t cell development. Genes & development 13, 1073–1078 (1999).

24. Jin, J. & Wang, W. Influential features PCA for high dimensional clustering. Ann. Stat. 44, 2323–2359, DOI: 10.1214/15-AOS1423 (2016).

25. Andrews, T. S. & Hemberg, M. M3Drop: dropout-based feature selection for scRNASeq. Bioinformatics DOI: 10.1093/bioinformatics/bty1044 (2018).

26. Jiang, L., Chen, H., Pinello, L. & Yuan, G.-C. GiniClust: detecting rare cell types from single-cell gene expression data with Gini index. Genome Biol. 17, 144, DOI: 10.1186/s13059-016-1010-4 (2016).

27. Wang, X., Park, J., Susztak, K., Zhang, N. R. & Li, M. Bulk tissue cell type deconvolution with multi-subject single-cell expression reference. Nat. Commun. 10, 380, DOI: 10.1038/s41467-018-08023-x (2019).

28. Kingma, D. P. & Welling, M. Auto-Encoding Variational Bayes (VAE, reparameterization trick). ICLR (2014). arXiv:1312.6114v10.

29. Van Dyk, D. A. Nesting EM algorithms for computational efficiency. Stat. Sin. 10, 203–225 (2000).

30. Schroth, G. P. et al. From single-cell to cell-pool transcriptomes: Stochasticity in gene expression and RNA splicing. Genome Res. 24, 496–510, DOI: 10.1101/gr.161034.113 (2013).

31. Strehl, A. & Ghosh, J. Cluster ensembles — a knowledge reuse framework for combining multiple partitions. J. Mach. Learn. Res. 3, 583–617, DOI: 10.1162/153244303321897735 (2003).

