## Supplementary materials for "SIMPLEs: a single-cell RNA sequencing imputation strategy preserving gene modules and cell clusters variation"

Zhirui Hu, Songpeng Zu

January 13, 2020

#### 1 Inference procedure for SIMPLE and SIMPLE-B

##### 1.1 Nesting Monte Carlo EM algorithm

EM algorithm is often used to estimate the parameters in the missing data problem or models with latent structure. However, in our model, the expectation of log-likelihood over all the latent variables cannot be computed directly, so we applied a nesting Monte Carlo EM algorithm (Van Dyk, 2000) that alternatively uses Gibbs sampling to impute missing data from its conditional distribution at current parameters, compute the expectation over the indicators ($Z$) and factors ($F$) conditioning on the imputed data ($X$) and update the parameters ($B$, $\Lambda$, $\Sigma$) by maximizing the approximated expectation of full log-likelihood. The nesting Monte Carlo EM algorithm has been used in the hierarchical probit model (Van Dyk, 2000) and Monte Carlo EM algorithm has been used for censored data (Wei and Tanner, 1990). However, our model is much more complicated than previous models.

The expectation of full log-likelihood can be written as $E(\log P(X, F, Z|\Theta')|Y, \Theta))$ where we used $\Theta$ to denote all the parameters $(\mu, B, \Lambda, \Sigma, \pi)$. In an EM algorithm, E-step is to compute the expectation above and M-step to maximize over $\Theta'$. However, the expectation cannot be readily computed in our case. Thus, we decomposed the expectation into two steps: $E(\log P(X, F, Z|\Theta')|Y, \Theta)) = E_X(E_{F,Z}(\log P(X, F, Z|\Theta')|X, \Theta)|Y, \Theta)$. In each iteration, we approximated the outer expectation over $X$ by Monte Carlo samples of missing data at current $\Theta^t$. Given L copies of the imputed data $X^l$, we used an EM algorithm to maximize the $\max_{\Theta'} E_X(\log P(X|\Theta')|Y, \Theta^t) \approx \max_{\Theta'} \frac{1}{L}\sum_l \log P(X^l|\Theta')$. For notation simplicity, we still use $Eh(X)$ to represent the empirical mean of $\frac{1}{L}\sum_l h(X^l)$. In the E-step, we compute the expectation of log-likelihood over indicators ($Z$) and factors ($F$) which needs the following terms: for each cell, we compute:

$$p(z_i = m|X) \propto \pi_m e^{-(\vec{x}_i - \vec{\mu}_m)^T (B\Lambda_m B^T + \Sigma_m)^{-1}(\vec{x}_i - \vec{\mu}_m)/2} \tag{1}$$

$$\begin{aligned} E(\vec{f}_i|z_i = m, X) &= (B^T \Sigma_m^{-1} B + \Lambda_m^{-1})^{-1} B^T \Sigma_m^{-1} (\vec{x}_i - \vec{\mu}_m) \\ Cov(\vec{f}_i|z_i = m, X) &= (B^T \Sigma_m^{-1} B + \Lambda_m^{-1})^{-1} \end{aligned} \tag{2}$$

In the M-step, we update $B,\ \Lambda,\ \Sigma$ and $\mu$ sequentially. For each gene $g$, we maximize $B_g$ by solving a linear regression with $L_1$ penalty:

$$arg \max_{\vec{B}_g} -\frac{1}{L}\sum_l E\left[\sum_{i,m} \frac{(x^l_{gi} - \mu_{gm} - \vec{B}_g{}^T \vec{f}_i)^2}{2\sigma^2_{gm}} I(z_i = m)\Big| X^l\right] - \frac{||\vec{B}_g||_1}{2\lambda}$$

which can be written as a penalized linear regression problem with the objective function: $\min_{\vec{B}_g} ||\tilde{y} - \widetilde{X}\vec{B}_g||_2^2 + \frac{L||\vec{B}_g||_1}{\lambda}$ where

$$\begin{aligned} \tilde{y}_{i,l,m} &= \frac{x^l_{gi} - \mu_{gm}}{\sigma_{gm}} \sqrt{p(z_i = m|X^l)} \\ \widetilde{X}_{i,l,m} &= \frac{E[\vec{f}_i I(z_i = m)|X^l]}{\sigma_{gm}\sqrt{p(z_i = m|X^l)}},\ i = 1, 2, ..., n,\ m = 1, 2, ..., M,\ l = 1, 2, ..., L \end{aligned} \tag{3}$$

and let $M = \sum_{i,m,l} \frac{Cov(f_i|z_i=m,X^l)}{\sigma^2_{gm}} \cdot p(z_i = m|X^l)$, a $K \times K$ matrix, then $\tilde{y}_j = 0,\ j = nLM + 1, .., nLM + K,\ \tilde{X}_{nLM+1,nLM+K} = M^{\frac{1}{2}}$ where $\tilde{X}_{nLM+1,nLM+K}$ is the $nLM + 1$ to $nLM + K$ rows of $\tilde{X}$. To maximize over $\Lambda$, we need to solve a non-linear function with linear constraint: for each $k$,

$$\max_{\lambda_{km}} -\frac{\sum_i E[f^2_{ik} I(z_i = m)]}{\lambda^2_{km}} - \log \lambda^2_{km} \cdot \sum_i p(z_i = m) \tag{4}$$

subject to $\sum_m \lambda^2_{km} = 1$. After updating $B$ and $\Lambda$, we update each $\sigma_{gm}$ by:

$$\max_{\sigma} -\frac{\sum_i E[(x_{gi} - \mu_{gm} - \vec{B}_g{}^T \vec{f}_i)^2 I(z_i = m)]}{2\sigma^2} - \frac{1}{2}\log \sigma^2 \cdot \sum_i p(z_i = m) - (\frac{\alpha}{2} + 1)\log \sigma^2 - \frac{\beta}{2\sigma^2} \tag{5}$$

The first term could be computed from the residual sum of squares of the regression problem above. The optimum value of $\sigma_{gm}$ has a closed-form solution by taking derivative of $\sigma$. Finally, we updated $\mu_m$ for each cluster by maximizing the log-likelihood integrating out $F$:

$$arg \max_{\vec{\mu}_m} -E\left[\sum_i (\vec{x}_i - \vec{\mu}_m)^T (B\Lambda_m B^T + \Sigma_m)^{-1} (\vec{x}_i - \vec{\mu}_m) I(z_i = m)\right] - \frac{||\vec{\mu}_m||^2}{\sigma_0^2}$$

then

$$\hat{\mu}_m = \left[\sum_i E[I(z_i = m)] + \frac{B\Lambda_m B^T + \Sigma_m}{\sigma_0^2}\right]^{-1} \sum_i E[\vec{x}_i I(z_i = m)] \tag{6}$$

in which the matrix inversion can be computed efficiently by Woodbury formula.

We used several steps of Gibbs sampling to obtain Monte Carlo samples of the missing values conditioning on the current parameters. At each iteration, the Gibbs sampler is initialized from the imputed data from last Monte Carlo EM iteration. For each cell $i$, we first sample cluster membership $z_i$ by (1) and the factors $\vec{f_i}$, then sample the values for zero entries conditioning on $\vec{f_i}$ and $z_i$. We sample $\vec{f_i}$ conditional on $z_i$ and others according to:

$$\vec{f_i}|z_i = m, \mu, B, \Lambda, \Sigma, \vec{x_i} \sim N(\vec{f_i}|(B^T\Sigma_m^{-1}B + \Lambda_m^{-1})^{-1}B^T\Sigma_m^{-1}(\vec{x_i} - \vec{\mu}_m), (B^T\Sigma_m^{-1}B + \Lambda_m^{-1})^{-1}) \tag{7}$$

Conditioning on $\vec{f_i}$ and $z_i$, the imputed values for each gene can be sampled independently. For each gene, we first sample an indicator for whether the zero entry is because of dropout and then sample $x_{gi}$:

$$\begin{aligned}
P(I_{gi} = 1|\vec{f_i}, z_i = m, \vec{B}_g, \mu_{gm}, \sigma_{gm}^2, y_{gi} = 0) &\propto (1 - p_g) \\
P(I_{gi} = 0|\vec{f_i}, z_i = m, \vec{B}_g, \mu_{gm}, \sigma_{gm}^2, y_{gi} = 0) &\propto p_g\Phi(-\frac{\mu_{gm} + \vec{B}_g^T\vec{f_i}}{\sigma_{gm}^2}) \\
x_{gi}|I_{gi} = 1, \mu_{gm}, \sigma_{gm}^2, y_{gi} = 0 &\sim N(x_{gi}|\mu_{gm} + \vec{B}_g^T\vec{f_i}, \sigma_{gm}^2) \\
x_{gi}|I_{gi} = 0, \mu_{gm}, \sigma_{gm}^2, y_{gi} = 0 &\propto N(x_{gi}|\mu_{gm} + \vec{B}_g^T\vec{f_i}, \sigma_{gm}^2)I(x_{gi} < 0)
\end{aligned} \tag{8}$$

$I_{gi}$ is an indicator for dropout event and $\Phi$ is the cdf of a standard normal distribution.

The overall algorithm shown in Algorithm 1 can output $\hat{B}$ which characterizes the gene-gene correlation within cluster, $\Lambda$ indicating which factor has larger effect in each cluster, and the cluster membership that takes account of the dropout events. We can impute the data by the posterior mean and use the imputed data for downstream analysis. If using the imputed data for differential gene expression analysis, one might also need to take account of the uncertainty, e.g. the posterior variances, of the imputed values.

#### 1.2 Initializing clustering membership

We found that using PCA for dimension reduction and then using (weighted) Kmeans for clustering can give a good initialization of cells cluster membership. Spectral clustering is commonly used for single cell RNASeq (e.g. Seurat (Butler et al., 2018), SC3 (Kiselev et al., 2017)). First, we did PCA on the standardized gene expression matrix and mapped each cell to the top PCs. To be more specific,

**Algorithm 1** Monte Carlo EM algorithm for imputation and clustering

**Input:** $\{y_{gi}\}$ log-normalized gene expression single cell RNASeq, $M$ number of clusters, $K$ number of latent factors, $p_{min}$ as a upper bound for dropout rate.
**Initialization:**

1: Initialize clustering membership by PCA then Kmeans.
2: Obtain $\hat{p}_g$ and initial values of $\mu_{gm}$, $\{x_{gi}\}$ for "high quality" genes by (9).
3: Nesting Monte Carlo EM algorithm for "high quality" genes:
4: Initialize $B \sim N(0, \frac{1}{\sqrt{K}})$, $\lambda_{km} = \frac{1}{\sqrt{M}}$, $\sigma_{gm} = 1$, $\pi_m = \frac{1}{M}$.
5: **for** iter = 1 to max iteration **do**
6:     E-step: compute expectations of $F$ and $Z$ by (2) and (1).
7:     M-step: update $B$ by (3), $\Lambda$ by (4), $\Sigma$ by (5), $\mu$ by (6).
8:     sample $Z$ by (1), $F$ by (7) and then sample new imputed values $x_{gi}$ by (8).
9: **end for**
10: Output sub-matrices of $\hat{B}$, $\hat{\Sigma}$, $\hat{\mu}$, $\hat{\Lambda}$ and the expected values of $F$ and $Z$ by (2) and (1) respectively.
11: Compute sub-matrices of $\hat{B}$, $\hat{\Sigma}$, $\hat{\mu}$ for "low quality" genes by (3) and (5), set $\hat{p}_g = p_{min}$ and impute the zero entries of these genes by (8).
12: **Estimation:** Repeat line 5 to 9 for all the genes and output $\hat{B}$, $\hat{\Sigma}$, $\hat{\Lambda}$, $\hat{\mu}$.
13: **Imputation:** Repeat line 8 fix the parameters at MLE to obtain the posterior distribution of imputed values.
14: **Output:** $\{x_{gi}\}$ imputed gene expression, $z_i$ clustering membership.

let $Y = UDV^T$ be the singular value decomposition of the gene expression matrix, where $U$ is a $G \times n$ orthogonal matrix, $D$ is a $n \times n$ diagonal matrix and $V$ is a $n \times n$ orthogonal matrix (in most cases $G > n$). We only projected the data matrix onto the top $K$ left singular vectors, then we did Kmeans clustering using the first $K$ columns of $VD$. Choosing the number of PCs should depend on the number of cells. We used 20∼50 PCs for 200∼700 cells in our examples. Next, we used Kmeans on the projected matrix, which is equivalent to a Gaussian mixture model assuming the same diagonal co-variance matrix for each cluster.

#### 1.3 Initializing imputation and factors

We initialized $B = 0$ and thus each gene is independent for cells in the same cluster. We treated genes with high or low number of zeros differently for initial imputation. For genes with many zero entries, it is hard to distinguish dropout from amplified component only based on the marginal distribution of gene expression. Thus, we first imputed zero entries and estimated $F$ as well as other parameters only using genes with few zero entries ("high quality" genes) and then projected the expression of other genes onto $F$. We found that estimating $F$ only using genes with few zero entries is better than using all the genes, since the number of latent factors are low and each factor loading is not so sparse that the factor can be only recovered by a few genes. On the other hand, initial imputation for genes with many zeros can be quite arbitrary and including these genes can drive the factors towards some local modes.

To initialize imputation, we first fitted a zero-inflated censored normal model of each "high quality" gene for cells of each type and obtained MLE of the mean $\hat{\mu}_g$, the variance $\hat{\sigma}_g^2$ and $\hat{p}_g$. Then, we impute missing values according to:

$$P(x_{gi}|\hat{\mu}_g, \hat{\sigma}_g^2, y_{gi} = 0) \propto N(x_{gi}|\hat{\mu}_g, \hat{\sigma}_g^2)I(x_{gi} < 0)\hat{p}_g + N(x_{gi}|\hat{\mu}_g, \hat{\sigma}_g^2)(1 - \hat{p}_g) \quad (9)$$

which is the same as (8) but setting $B = 0$. Then we used the nesting Monte Carlo EM algorithm to obtain sub-matrices of $\hat{B}$, $\hat{\Sigma}$ for "high quality" genes, $\hat{\Lambda}$ as well as the expected values of $F$ and $Z$ by (2) and (1) respectively. Based on these expected values, we computed sub-matrices of $\hat{B}$, $\hat{\Sigma}$ for genes with many zeros by (3) and (5) and imputed the zero entries of these genes by (8) using the expected values of $F$ and samples of $Z$ obtained from "high quality" genes. In the real example, we set a upper bound on dropout rate which can be suggested by an experimentally estimate of the dropout rate in the single cell RNASeq (Schroth et al., 2013) or empirically from overall dropout rate estimates in the data set. For "low quality" genes, we set the dropout rate to be the upper bound otherwise the data can be unduly imputed.

#### 1.4 Clustering stability

If we have multiple copies of imputed expression matrix, we did the clustering procedure for each imputation and summarized the frequency that each pair of cell is in the same cluster into a consensus clustering matrix. Based on this matrix, a further hierarchical clustering could be conducted to get a more stable clustering result(Kiselev et al., 2017). We further defined the consensus score for each pair of the cell as $s_{ij} = \min\{p_{ij}, 1 - p_{ij}\}$, where $p_{ij}$ is the frequency that cell $i$ and $j$ are in the same cluster. Then, we assigned an uncertainty score for each cell averaging over the consensus score above, i.e. $s_i = avg_j(s_{ij})$.

#### 1.5 Model checking

In each cell type, we assumed a zero-inflated censored Gaussian distribution for each gene marginally. To validate our assumption on zero component, we compared the percentage of zeros observed for each gene with the expected percentage of zeros under a negative-censored Gaussian distribution. In detail, we fitted a truncated normal distribution for each gene only using positive counts and computed the p-values for the observed number of zeros using a binomial test. For the "hESC cell type data", we identified 8% to 16% genes with FDR < 0.05 (around 1000 to 2000 genes) in each of the 7 cell types, which had fewer zeros than predicted by the fitted Gaussian distribution using only positive counts. These genes in general had large fitted variances, so that the expected probability of zero counts is high. The large variances might be due to heterogeneity within a cell type, especially for DECs, which can be modeled by SIMPLEs. Thus, the number above might over-estimate the actual number of genes that does not fit the zero-inflated censored Gaussian distribution well. Compared with "hESC cell type data", mouse embryonic data set has more dropouts. For mouse embryonic data set, we only identified 2% to 7% genes with FDR $< 0.05$ (around 200 to 800 genes) in each of the 6 cell types, which had fewer zeros than predicted by the fitted Gaussian distribution using only positive counts. Similarly, blastocytes have the largest number of genes with fewer zeros than expected, as gene expression in blastocytes has large variation as shown in the analysis in the main text.

#### 2 Supplementary Figures

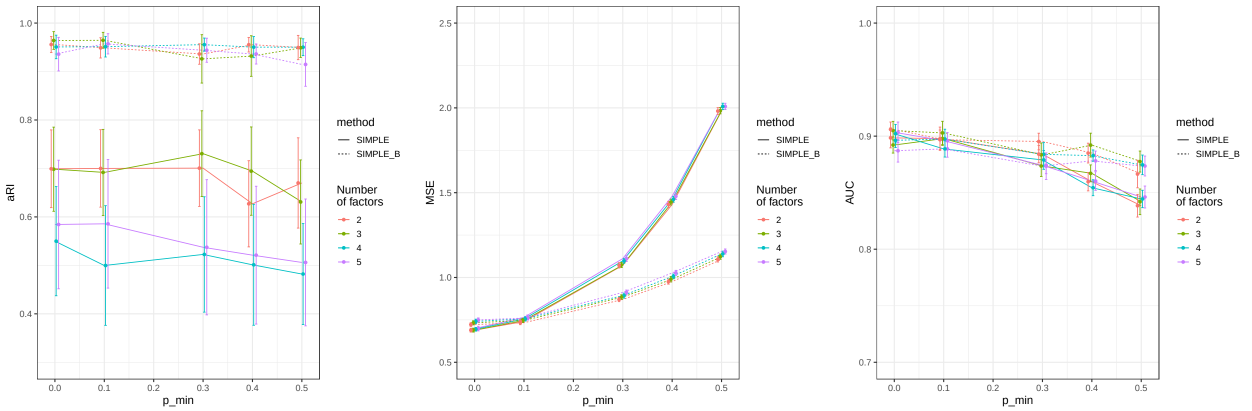


Supplemenary Fig. 1: Varying parameters for SIMPLE and SIMPLE-B for simulating independent gene expression with high dropout rate. X axis: minimum of prior amplified rate for SIMPLE and SIMPLE-B respectively; Y axis: from left to right are adjusted rand index, mean squared error of imputed values and AUC for identifying marker genes respectively. The error bar shows the standard deviation over 10 repetitions. Colors of the curves represent different number of factors used in the program and line styles represent either SIMPLE or SIMPLE-B. In this experiment, we fixed the number of high quality genes at 260.

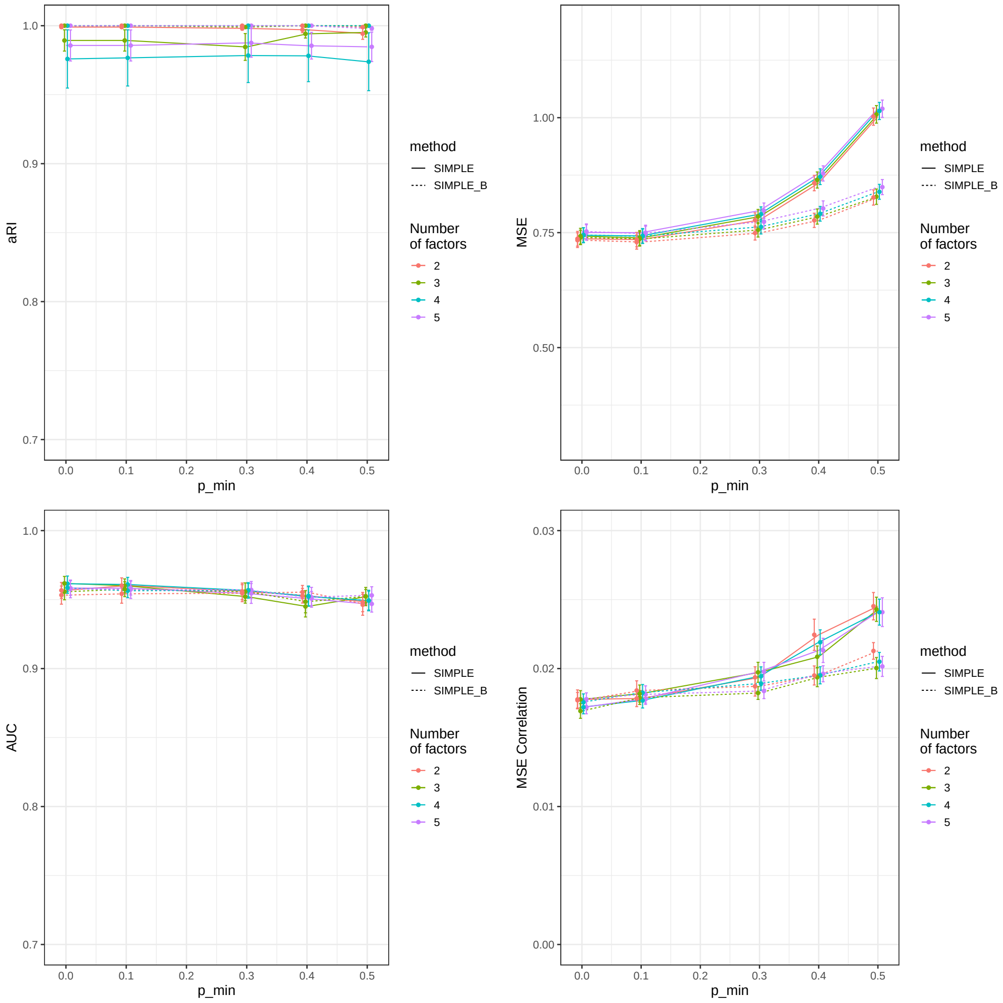


Supplemenary Fig. 2: Varying parameters for SIMPLE and SIMPLE-B for simulating correlated gene expression with low dropout rate. X axis: minimum or prior amplified rate for SIMPLE and SIMPLE-B respectively; Y axis: adjusted rand index, mean squared error of imputed values, AUC for identifying marker genes and mean squared error of estimating nonzero correlation within each cluster. The error bar shows the standard deviation over 10 repetitions. Colors of the curves represent different number of factors used in the program and line styles represent either SIMPLE or SIMPLE-B. In this experiment, we fixed the number of high quality genes at 700.

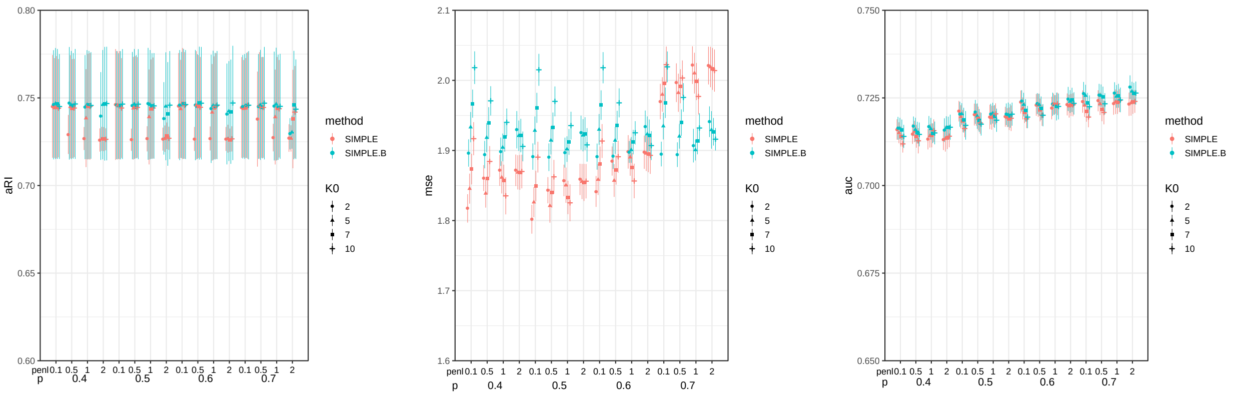


Supplemenary Fig. 3: Varying parameters for SIMPLE and SIMPLE-B for simulating dropouts based on hESC cell types data set. X axis: "p" is the minimum or the prior amplified rate for SIMPLE and SIMPLE-B respectively and "penl" is the penalization parameter of the prior distribution of weight matrix; Y axis: adjusted rand index, mean squared error of imputed values, and AUC for identifying marker genes. The error bar shows the 2*standard deviation over 10 repetitions. The shape represents different number of factors and the color represents either SIMPLE or SIMPLE-B. In this experiment, we fixed the number of high quality genes at 1500.

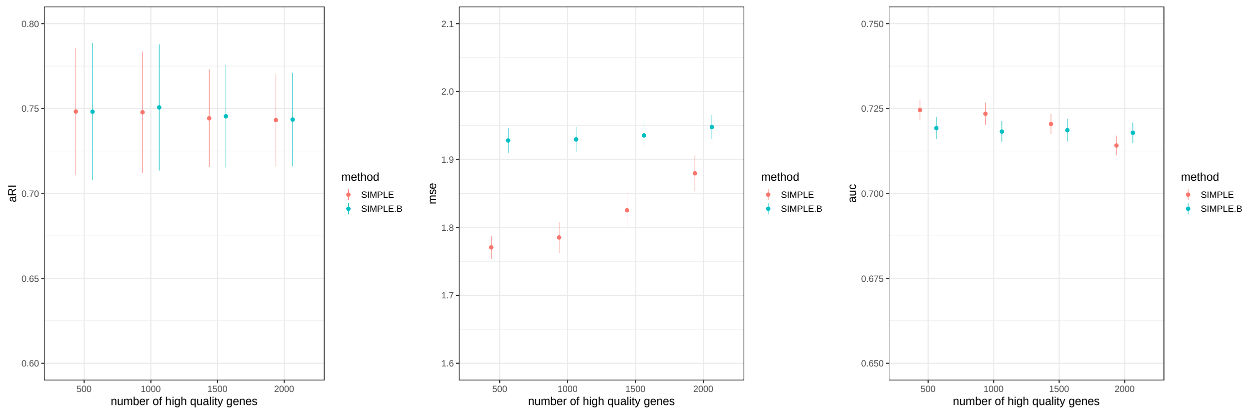


Supplemenary Fig. 4: Varying parameters for SIMPLE and SIMPLE-B for simulating dropouts based on hESC cell types data set. X axis: the number of high quality genes (with fewer zero entries) used for initialization; Y axis: adjusted rand index, mean squared error of imputed values, and AUC for identifying marker genes. The error bar shows the 2*standard deviation over 10 repetitions. The color represents either SIMPLE or SIMPLE-B. In this experiment, we fixed the prior of amplified rate ($p_m$) at 0.5, number of factors at 10 and penalty parameter at 1. For SIMPLE, the performance is better if we only use 500 genes with fewest zero entries for initialization, indicating that the cell type information can be obtained using only a few genes.

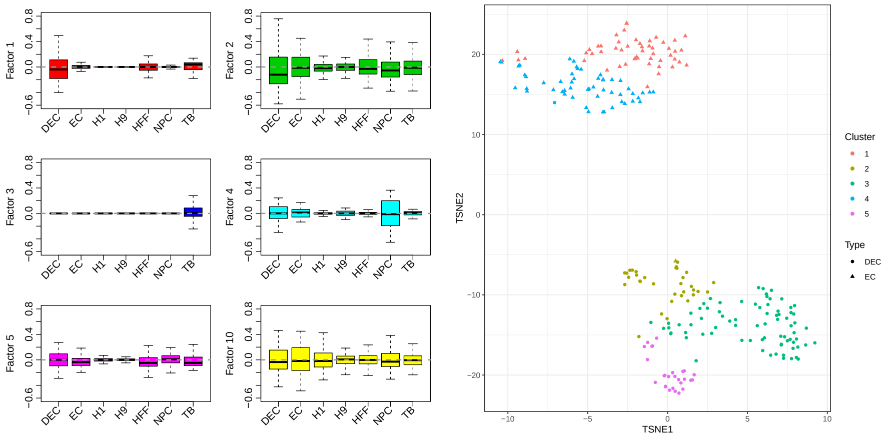


(a) Distribution of cells' latent factors (b) Subtypes of DEC and EC

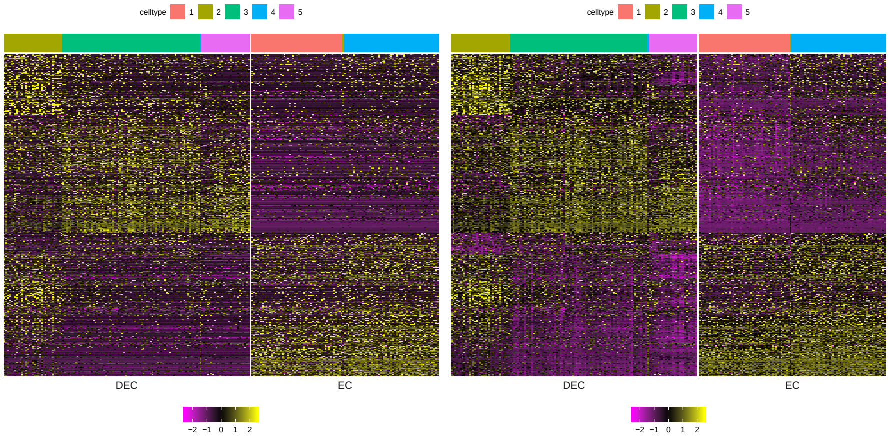


(c) Original data (d) Imputed by SIMPLE

Supplemenary Fig. 5: Subtypes of DECs and ECs in the hESC cell types data set. (a) Distribution of cells' latent factors in each cell type. We set $K = 10$ in our algorithm but only 6 of them having nonzero factor loadings. In each subfigure, the boxplot shows the quantiles of $E(f_{ik}) \cdot \overline{|B_k|} = E(E(f_{ik}|z_i)) \cdot \overline{|B_k|}$ over cell $i$ in each cell type for each k. $\overline{|B_k|}$ is the average factor loadings over nonzero entries of factor k, since $B_k$ might have different scales. (b) t-SNE plot using 1000 genes with large absolute values of coefficients of factor 1 ($B_1$) and expressed in at least 10% DECs and ECs. We extracted DECs and ECs from the imputed gene expression matrix, and clustered DECs and ECs into 5 subtypes using Kmeans based on these genes. Each dot is a cell and its color indicates the clustering result and shape indicates the true cell label. Results did not change much varying the number of selected genes. (c-d) Gene expression heatmap using (c) original or (d) imputed data in DECs and ECs. For clarity, only shows top 500 genes with largest absolute values of coefficients among the 1000 genes for clustering. Each row is a gene and each column is a cell first ordered by true cell label and then ordered by clusters. The cluster labels are shown by the colorbar above the heatmap. The color indicates the z-score (gene-wise centered and scaled).

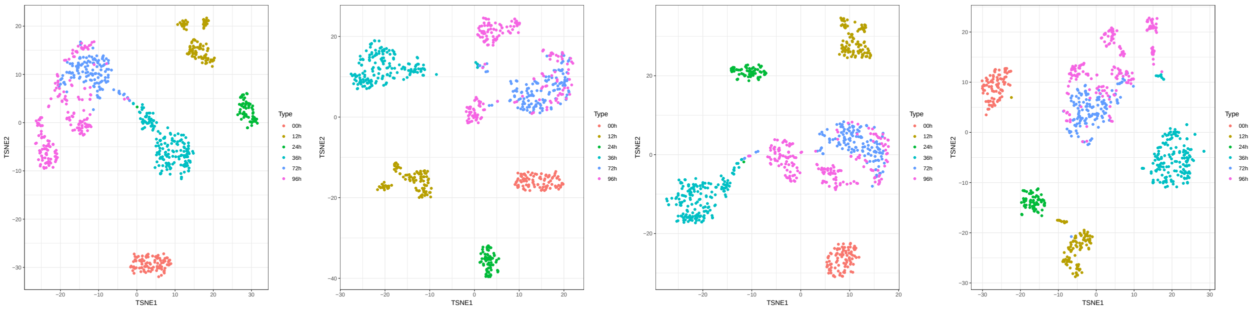


(a) Original data (b) SIMPLE (c) SIMPLE-B (d) scImpute

Supplemenary Fig. 6: Visualizing the original data or imputed data by different methods using tSNE for hESC time course data set. Each point is a cell colored by the time point. For clustering, the adjusted rand index are 0.74 using original data, 0.78 and 0.75 using SIMPLE and SIMPLE-B respectively, and 0.73 using scImpute.

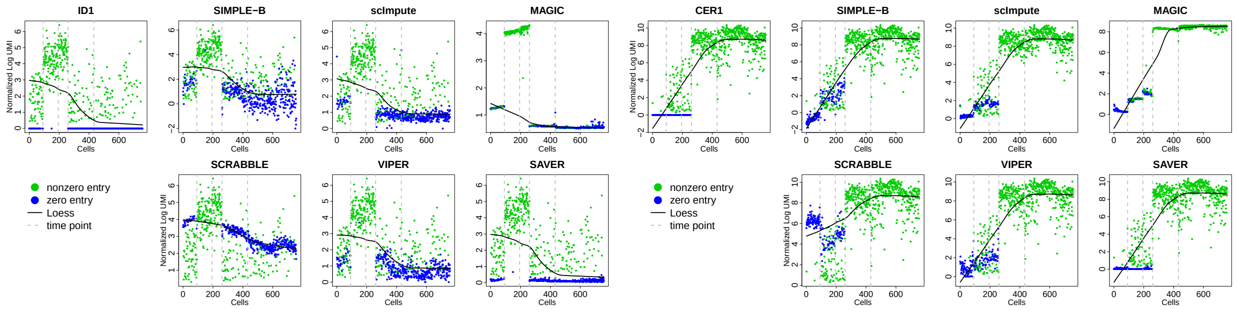


(a) ID1 (b) CER1

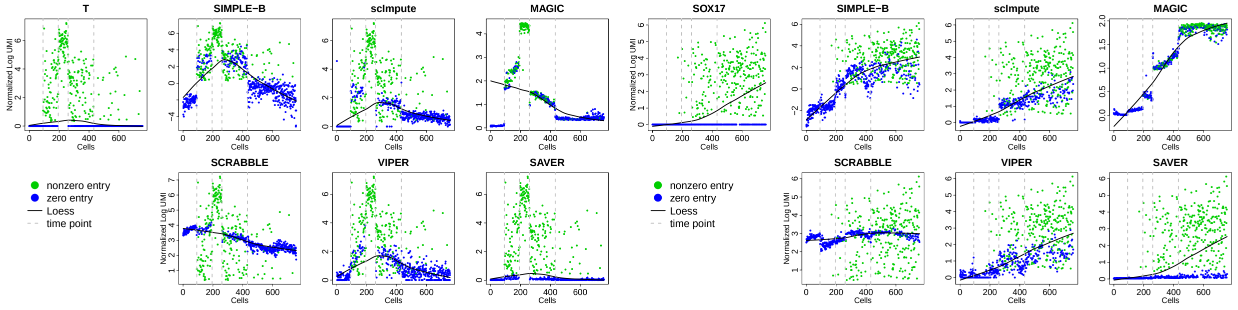


(c) T (d) SOX17

Supplemenary Fig. 7: Examples of imputed marker gene expression along cell developmental pseudo-time. Each dot is the gene expression level in a cell. Green dot: nonzero expression. Blue dot: zero expression which is imputed by different methods. The smoothed curve is computed loess using either the original or imputed data. Cells are first ordered by time label, then within each time, are ordered by pseudo-time output by Monocle. The dashed lines are the boundary of 0h, 12h, 24h, 36h. We treated cells at 72h and 96h as one group and ordered them by pseudo time. (a) ID1, highly expressed in the cells at early stage responding to the chemical treatment to induce differentiation; (b) CER1 is a early DE-specific genes which is up-regulated at 36 h of differentiation; (c) T, a mesendodorm marker, expressed when cell transits from a multi-potent state towards several more differentiated lineages; (d) SOX17, key DE markers activated as early as 36h.

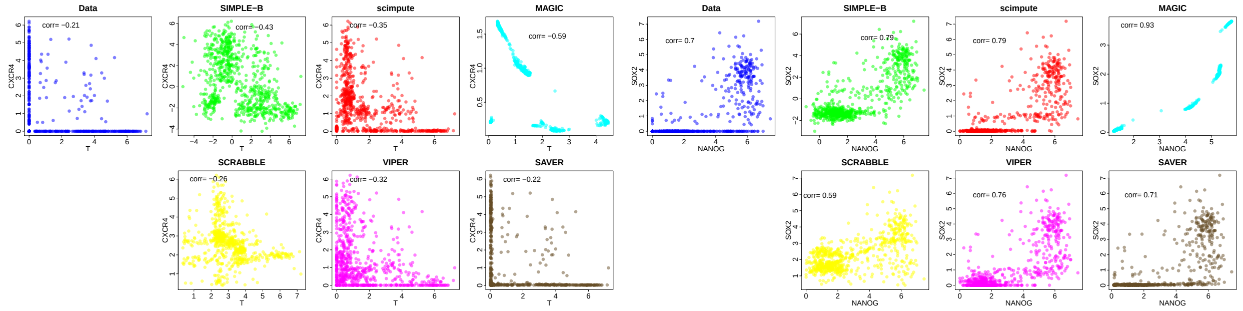


(a) T and CXCR4 (b) NANOG and SOX2

Supplemenary Fig. 8: Examples of correlated gene pairs from hESC time course data. Original or imputed expression of (A) T and CXCR4 (B) NANOG and SOX2 by different methods. Each dot represents a cell and the correlations between these two genes are shown in the left corner of each sub-figure.

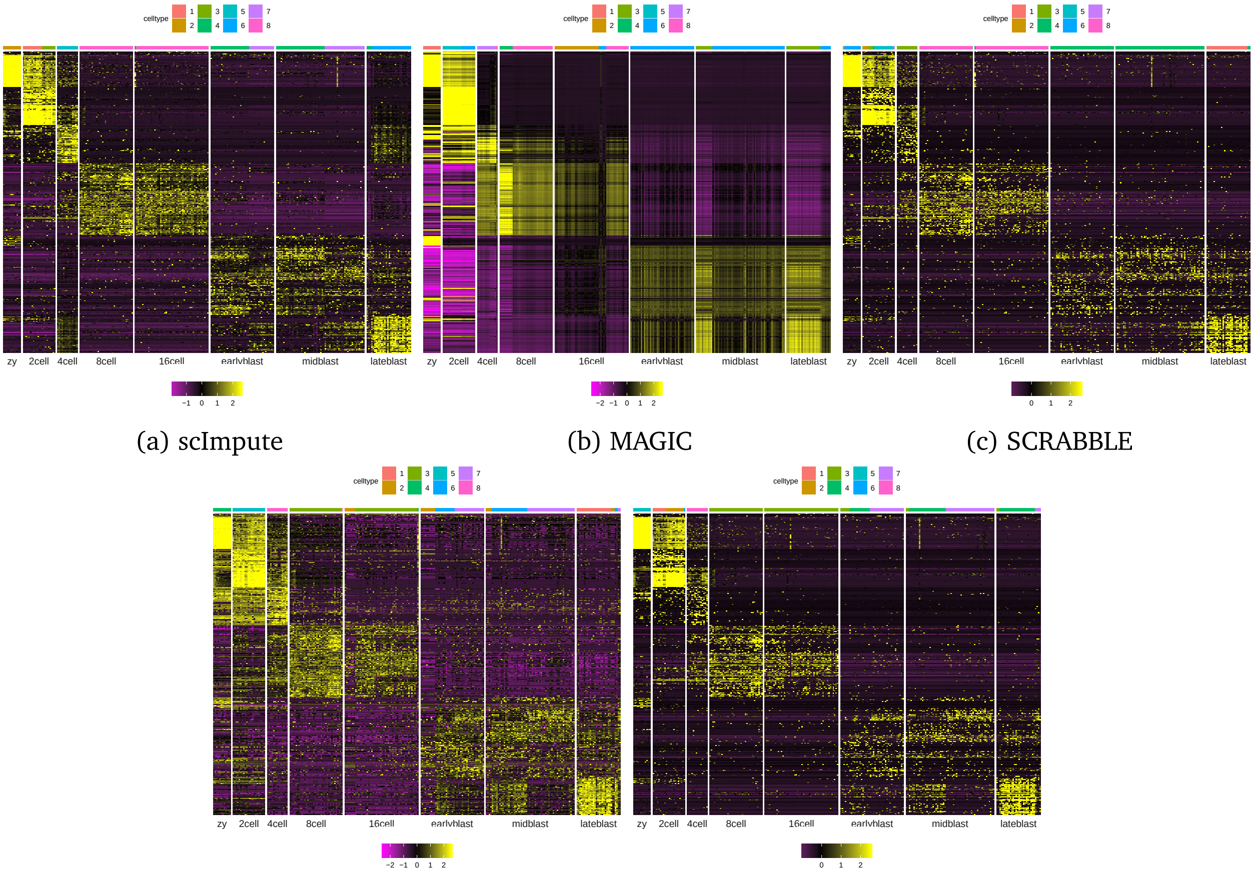


(a) scImpute (b) MAGIC (c) SCRABBLE

(d) VIPER (e) SAVER

Supplemenary Fig. 9: Imputed gene expression by different methods for mouse embryos data set. Each row is a marker gene and each column is a cell ordered subtypes. The purple to yellow color indicates the z-score range (centered and scaled by gene). The color bar above the heatmap shows the clustering result by each imputed matrix. The orders of the genes are the same as in Figure 5, but the cells within each subtype are ordered by clusters obtained using each imputation method. The differential expressed genes were identified by Wilcoxon test. For scImpute, we set the number of clusters equal to 6 which is the number of major cell types.

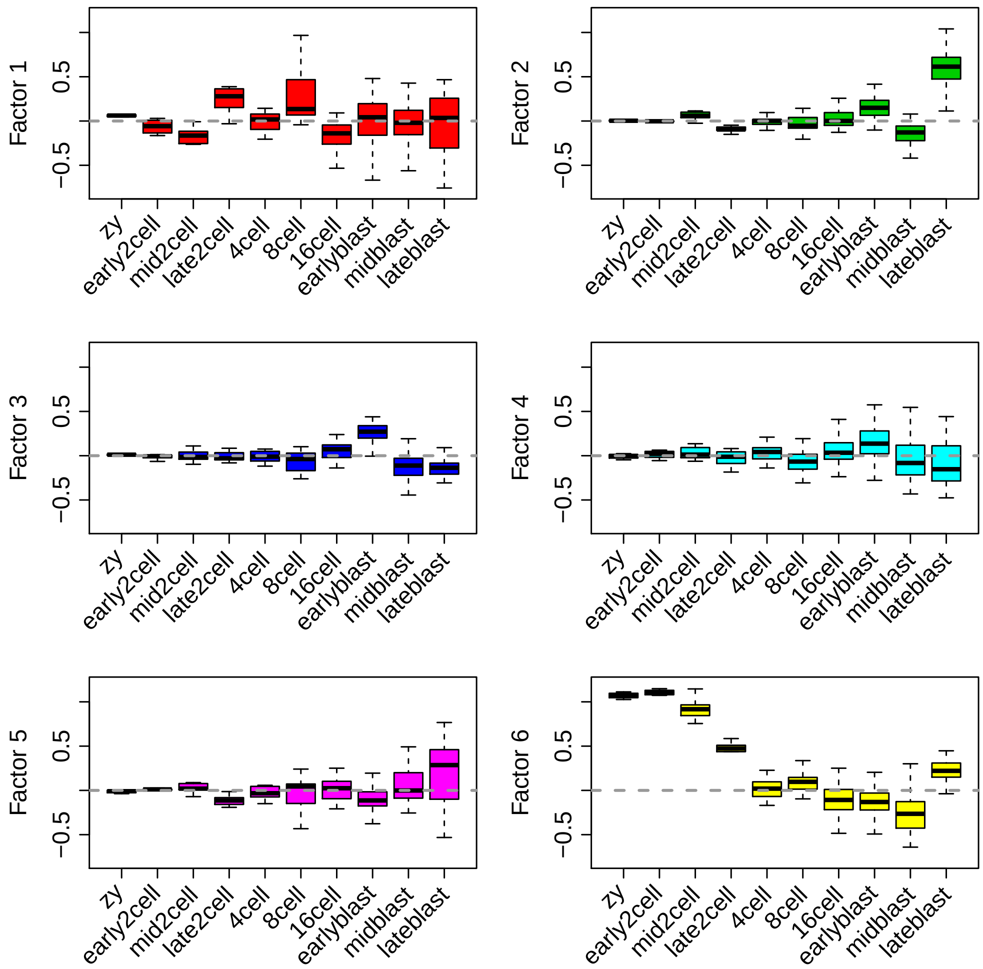


Supplemenary Fig. 10: Distribution of cells' latent factors in each subtype of mouse embryos. We only showed 6 out of 8 factors whose distributions are different among subtypes. In each subfigure, the boxplot shows the quantiles of $E(f_{ik}) \cdot \overline{|B_k|} = E(E(f_{ik}|z_i)) \cdot \overline{|B_k|}$ over cell $i$ in each cell type for each k. $\overline{|B_k|}$ is the average factor loadings over nonzero entries for factor k, since $B_k$ might have different scales.

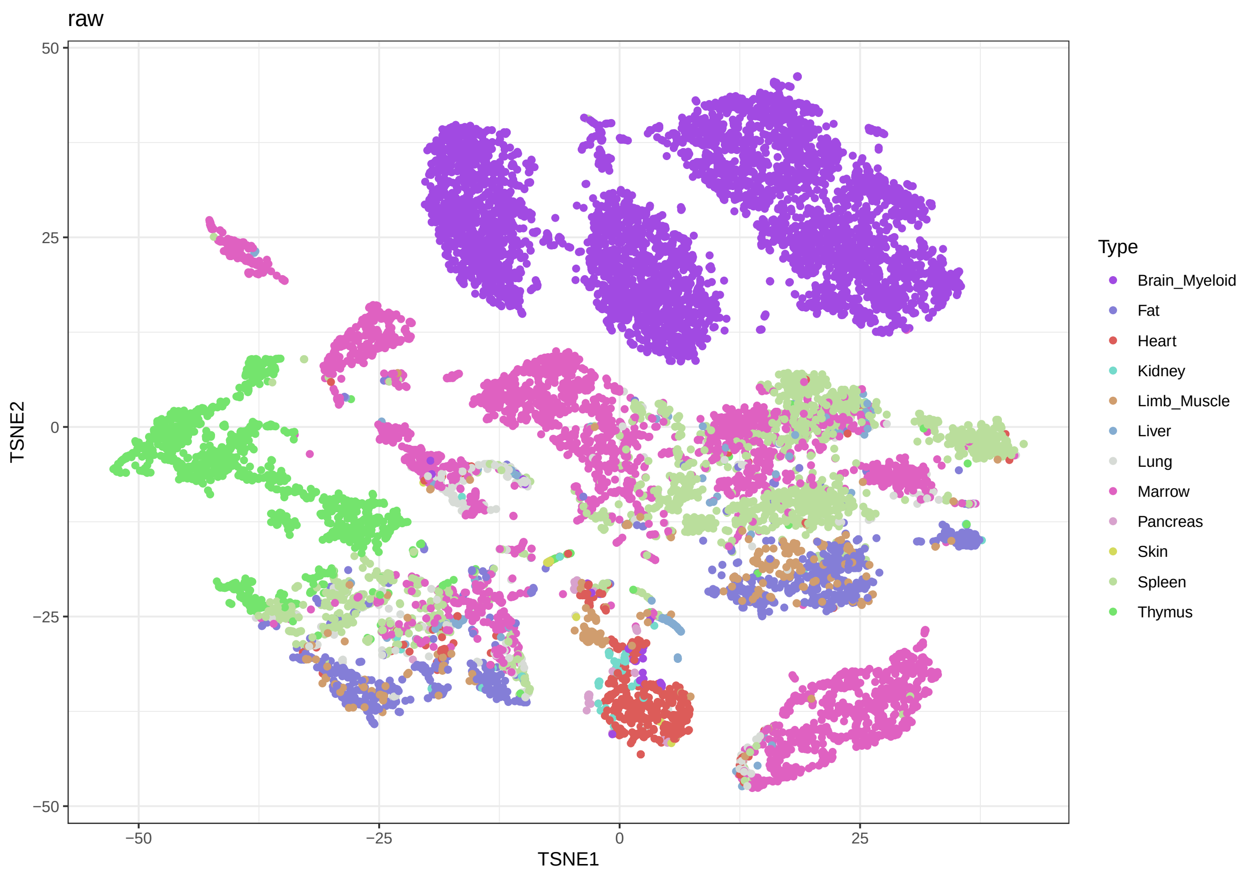


Supplemenary Fig. 11: T-SNE based data visualization of the immune cells using the original data. Each dot represents a cell and is colored by tissue.

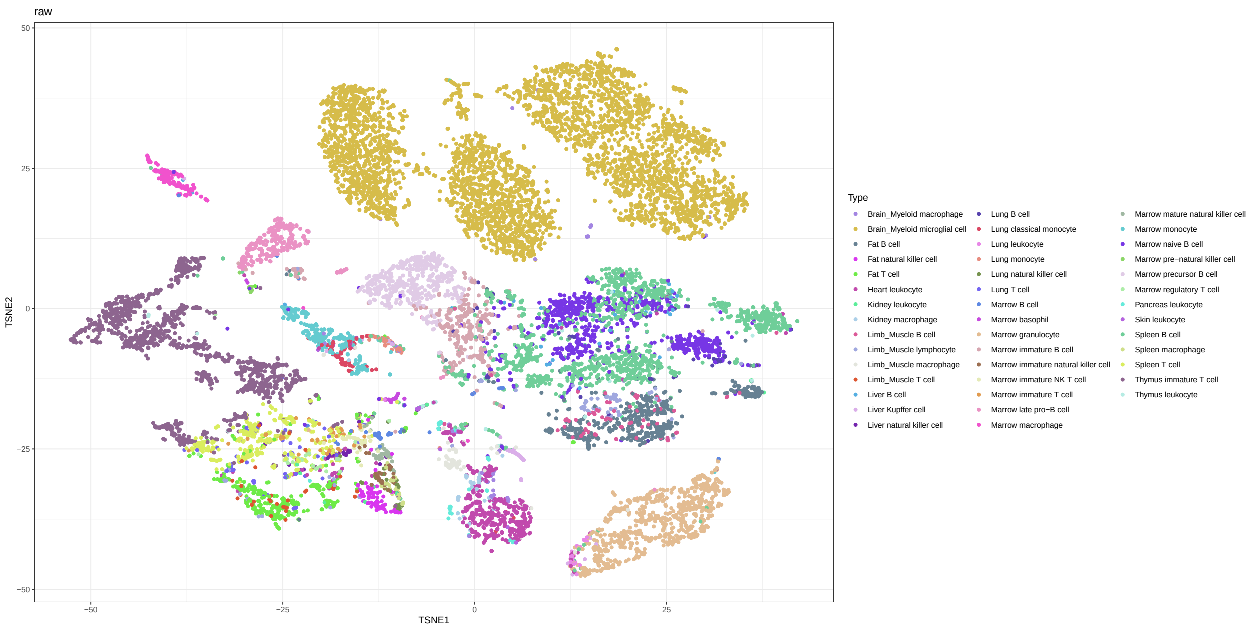


Supplemenary Fig. 12: T-SNE based data visualization of the immune cells using the original data. Each dot represents a cell and is colored by both tissue and cell type.

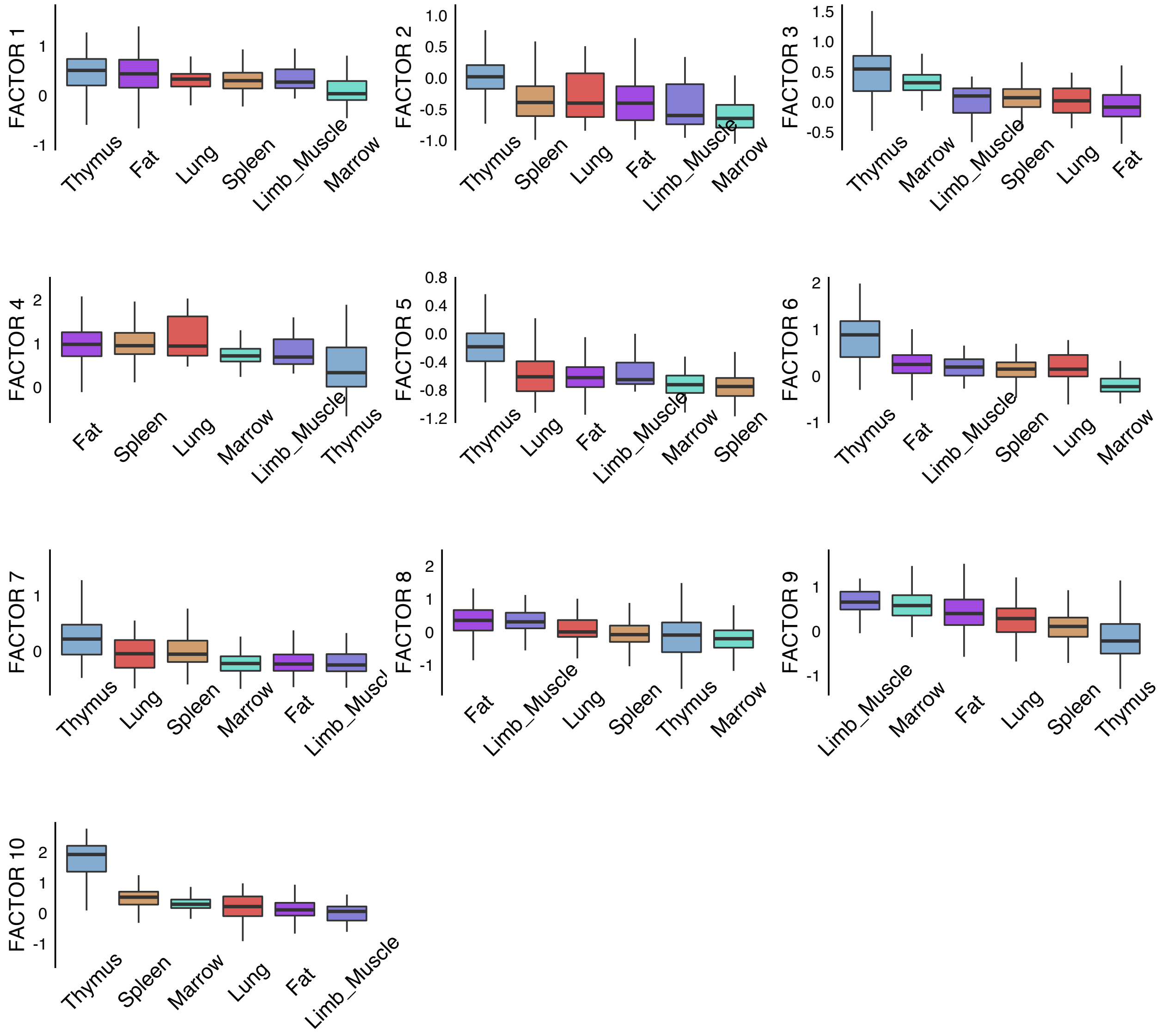


Supplemenary Fig. 13: Distribution of all the latent factors learned by SIMPLE for each tissue. This boxplot only includes T cells.

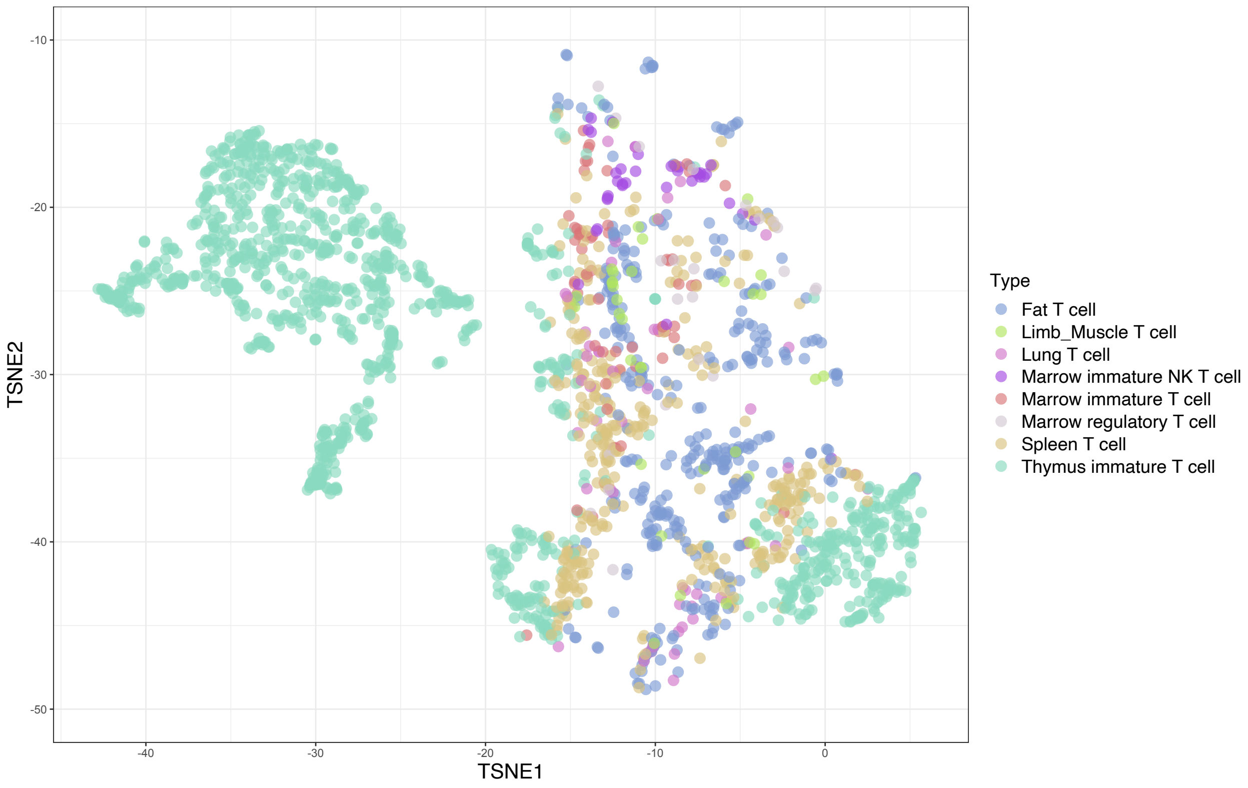


Supplemenary Fig. 14: T-SNE based data visualization of the T cells after imputation by SIMPLE. Each dot represents a cell and is colored by both tissue and cell type.

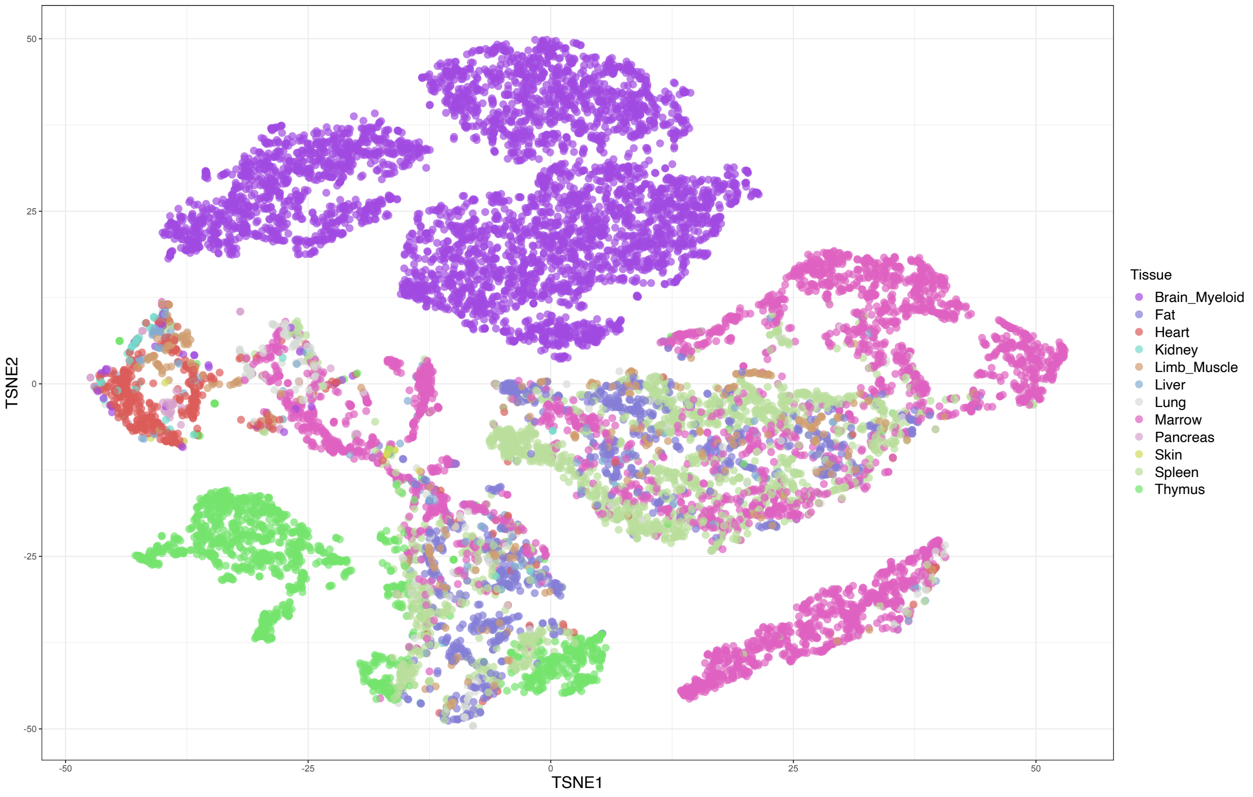


Supplemenary Fig. 15: T-SNE based data visualization of the immune cells after imputation by SIMPLE. Each dot represents a cell and is colored by both tissue type.

### References

Butler, Andrew, Paul Hoffman, Peter Smibert, Efthymia Papalexi and Rahul Satija. 2018. "Integrating single-cell transcriptomic data across different conditions, technologies, and species." *Nat. Biotechnol.* .

Kiselev, Vladimir Yu, Kristina Kirschner, Michael T. Schaub, Tallulah Andrews, Andrew Yiu, Tamir Chandra, Kedar N. Natarajan, Wolf Reik, Mauricio Barahona, Anthony R. Green and Martin Hemberg. 2017. "SC3: Consensus clustering of single-cell RNA-seq data." *Nat. Methods* 14:483–486.

Schroth, G P, J Gertz, R M Myers, B A Williams, K McCue, G K Marinov and B J Wold. 2013. "From single-cell to cell-pool transcriptomes: Stochasticity in gene expression and RNA splicing." *Genome Res.* 24(3):496–510.

Van Dyk, David A. 2000. "Nesting EM algorithms for computational efficiency." *Stat. Sin.* 10(1):203–225.

Wei, G.C.G. and M.A. Tanner. 1990. "A Monte Carlo Implementation of the EM Algorithm and the Poor Man's Data Augmentation Algorithms." *J. Am. Stat. Assoc.* 85(411):699–704.
